# Stress induces dynamic, cytotoxicity-antagonizing TDP-43 nuclear bodies via paraspeckle lncRNA *NEAT1*-mediated liquid-liquid phase separation

**DOI:** 10.1101/802058

**Authors:** Chen Wang, Yongjia Duan, Gang Duan, Qiangqiang Wang, Kai Zhang, Xue Deng, Beituo Qian, Jinge Gu, Zhiwei Ma, Shuang Zhang, Lin Guo, Cong Liu, Yanshan Fang

## Abstract

**Graphic Abstract:** 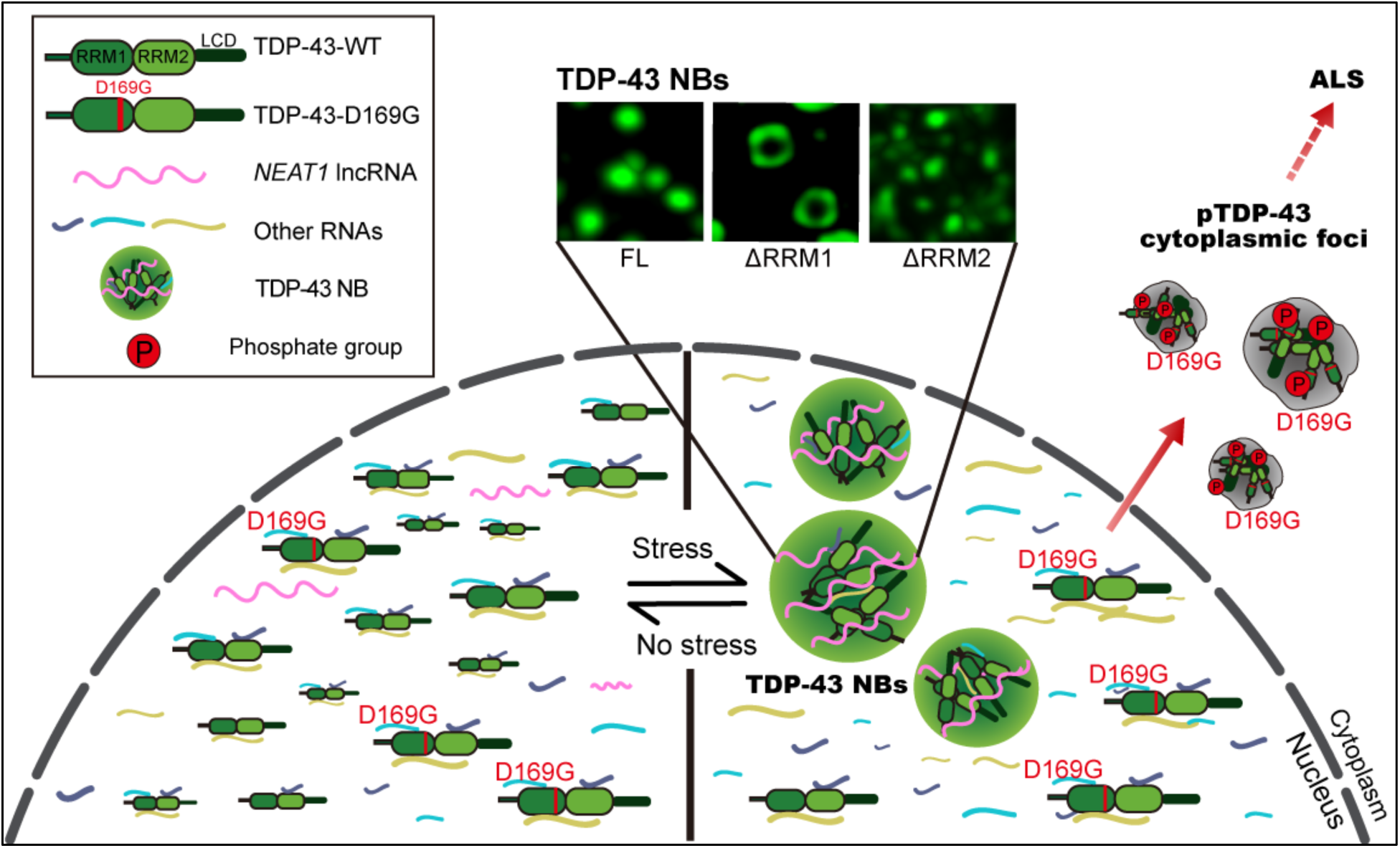

**Highlights:** *(Up to four bullet points. The length of each highlight cannot exceed 85 characters, including spaces)*

- Stress induces phase-separated TDP-43 NBs to alleviate cytotoxicity
- The two RRMs interact with different RNAs and act distinctly in the assembly of TDP-43 NBs
- LncRNA *NEAT1* promotes TDP-43 LLPS and is upregulated in stressed neurons
- The ALS-causing D169G mutation is NB-defective and forms pTDP-43 cytoplasmic foci

**Summary:** Despite the prominent role of TDP-43 in neurodegeneration, its physiological and pathological functions are not fully understood. Here, we report an unexpected function of TDP-43 in the formation of dynamic, reversible, liquid droplet-like nuclear bodies (NBs) in response to stress. Formation of NBs alleviates TDP-43-mediated cytotoxicity in mammalian cells and fly neurons. Super-resolution microscopy reveals a “core-shell” organization of TDP-43 NBs, antagonistically maintained by the two RRMs. TDP-43 NBs are partially colocalized with nuclear paraspeckles, whose scaffolding lncRNA *NEAT1* is dramatically upregulated in stressed neurons. Moreover, increase of *NEAT1* promotes TDP-43 liquid-liquid phase separation (LLPS) *in vitro*. Finally, we uncover that the ALS-associated mutation D169G impairs the *NEAT1*-mediated TDP-43 LLPS and NB assembly, causing excessive cytoplasmic translocation of TDP-43 to form stress granules that become phosphorylated TDP-43 cytoplasmic foci upon prolonged stress. Together, our findings suggest a stress-mitigating role and mechanism of TDP-43 NBs, whose dysfunction may be involved in ALS pathogenesis.

## Introduction

Amyotrophic lateral sclerosis (ALS) is a progressive motor neuron disease characterized by the degeneration of motor neurons in the brain and spinal cord, which leads to fatal paralysis generally within 2–5 years after diagnosis (Taylor et al., 2016; van Es et al., 2017). Missense mutations in the gene *TARDBP* encoding the trans-activation response element (TAR) DNA-binding protein 43 (TDP-43) predispose to familial ALS, and cytoplasmic protein inclusions containing pathologically phosphorylated and ubiquitinated TDP-43 in motor neurons are a pathological hallmark of ALS (Neumann et al., 2006; Kabashi et al., 2008). TDP-43 is a nuclear protein but can shuttle between the nucleus and the cytoplasm. It has a nuclear localization signal (NLS), a nuclear export signal (NES), two canonical RNA recognition motifs (RRMs) that bind to nucleic acids, and a C-terminal low complexity domain (LCD) that mediates protein-protein interactions and is enriched of disease-associated mutations (Kabashi et al., 2008; Lee et al., 2012). TDP-43 is engaged in a variety of ribonucleoprotein (RNP) complexes and plays an important role in RNA processing and homeostasis (Jovičić et al., 2016; Ratti and BuRatti, 2016; Ederle and Dormann, 2017; Fahrenkrog and Harel, 2018; Zhao et al., 2018). In addition, TDP-43 is known to participate in cytoplasmic stress granules (SGs), which may undergo aberrant phase transition and promote the formation of solid protein aggregates in diseased conditions (Li et al., 2013; Ramaswami et al., 2013; Murakami et al., 2015; Patel et al., 2015).

Nuclear bodies (NBs) are dynamic, membraneless nuclear structures that concentrate specific nuclear proteins and RNAs and play an important role in maintaining nuclear homeostasis and RNA processing (Stanek and Fox, 2017; Wegener and Müller-McNicoll, 2018). Increasing evidence suggests the association of TDP-43 with a variety of NBs, such as paraspeckles (Dammer et al., 2012; Nishimoto et al., 2013; West et al., 2016), gemini of coiled bodies (GEMs) (Ishihara et al., 2013), nuclear stress bodies (Udan-Johns et al., 2014), interleukin-6 and -10 splicing activating compartment bodies (InSAC bodies) (Lee et al., 2015), and omega speckles (Piccolo et al., 2018). Notably, in the nucleus of spinal motor neurons of sporadic ALS patients, TDP-43 is found colocalized with paraspeckles and the occurrence of paraspeckles is specifically increased in the early phase of the disease (Nishimoto et al., 2013). Excessive formation of dysfunctional paraspeckles is also observed in motor neurons of ALS-FUS patients (An et al., 2019). Paraspeckles are a class of subnuclear RNP granules formed by the scaffolding lncRNA *NEAT1* associated with splicing factor proline-glutamine rich (SFPQ), P54NRB/NONO and other paraspeckle proteins, which are engaged in regulating cellular functions through nuclear retention of mRNAs and proteins (Bond and Fox, 2009; Fox and Lamond, 2010; Nakagawa et al., 2018). Although TDP-43 is present in NBs such as paraspeckles that sometimes accompany neurodegeneration, our understanding about the function of TDP-43 in NBs and the involvement of NBs in ALS pathogenesis is still rudimentary.

Recent studies indicate that liquid-liquid phase separation (LLPS) of RNA-binding protein (RBPs) drives the assembly of liquid droplet (LD)-like, membraneless RNP granules in the cytoplasm and nucleoplasm (Lin et al., 2015; Feric et al., 2016; Banani et al., 2017; Uversky, 2017; Nakagawa et al., 2018; Fox et al., 2018). Several ALS-related RBPs including TDP-43, FUS, hnRNP A1, hnRNP A2 and TIA1 are shown to phase separate *in vitro* (Hyman et al., 2014; Molliex et al., 2015; Patel et al., 2015; Schmidt and Rohatgi, 2016; Mackenzie et al., 2017; Ryan et al., 2018; Wang et al., 2018). The intrinsically disordered LCD domains of the RBPs are thought to mediate the LLPS (Hennig et al., 2015; Molliex et al., 2015; Xiang et al., 2015; Murray et al., 2017; Uversky, 2017) and posttranslational protein modifications play an important role in determining the biophysical and biological properties of the phase behavior of the RBPs (Brady et al., 2011; Cohen et al., 2015; Li et al., 2017; Hofweber et al., 2018; Luo et al., 2018; McGurk et al., 2018; Qamar et al., 2018; Ryan et al., 2018; DuAn et al., 2019). In addition to the LCD, recent studies reveal that RNA is of vital importance in regulating the LLPS (Dominguez et al., 2018; Namkoong et al., 2018; Fox et al., 2018), which can either suppress or promote phase separation depending on the contents and concentrations of the RNAs (Shevtsov and Dundr, 2011; Lin et al., 2015; Maharana et al., 2018; Mann et al., 2019).

In this study, we find that various cellular stresses induce TDP-43 to form distinct, highly dynamic and reversible NBs, which not only attenuate the cytotoxicity of TDP-43 in mammalian cells but also ameliorate neurodegeneration and behavioral deficits in a *Drosophila* model of ALS. Further investigation with super-resolution microscopy uncovers the distinct functions of the two RRMs in maintaining the morphology and size of TDP-43 NBs. Moreover, we reveal a crucial role of the paraspeckle scaffolding long non-coding RNA (lncRNA) nuclear-enriched abundant transcript 1 (*NEAT1*) in promoting TDP-43 LLPS, which is compromised by the ALS-causing mutation D169G and causes a defect in TDP-43 NB formation. The NB-forming defective D169G mutant is prone to accumulate disease hallmarked, hyperphosphorylated TDP-43 foci in the cytoplasm with prolonged stress. Together, our findings propose that stress induces the assembly of TDP-43 NBs in the generally “suppressive” nucleoplasm environment via upregulation of lncRNA *NEAT1* which promotes TDP-43 LLPS, and the defective stress-mitigating TDP-43 NBs may contribute to ALS pathogenesis.

## Results

### Arsenic stress induces dynamic and reversible TDP-43 NBs

TDP-43 protein is predominantly localized to the nucleus but can shuttle between the nucleus and the cytoplasm. In response to cellular stress, TDP-43 is recruited to cytoplasmic SGs (Li et al., 2013). Interestingly, in an earlier related work of our group (DuAn et al., 2019), we noticed that although arsenite, a commonly used reagent to raise cellular stress (Bernstam and Nriagu, 2000), induced TDP-43^+^ cytoplasmic SGs, the majority of TDP-43 signal remained in the nucleus. More than that, in a higher percentage of the cells, nuclear TDP-43 lost the normal diffused pattern and instead exhibited a distinct granular appearance (Figures 1A and 1B, and Movie S1). Similarly, endogenous TDP-43 also formed granules in the nucleus upon arsenic stress (Figures 1C and 1D), with the size ranging from 0.01 to 0.67 µm^2^ and the median average of 0.019 µm^2^ (Figures 1E).

**Figure 1.**
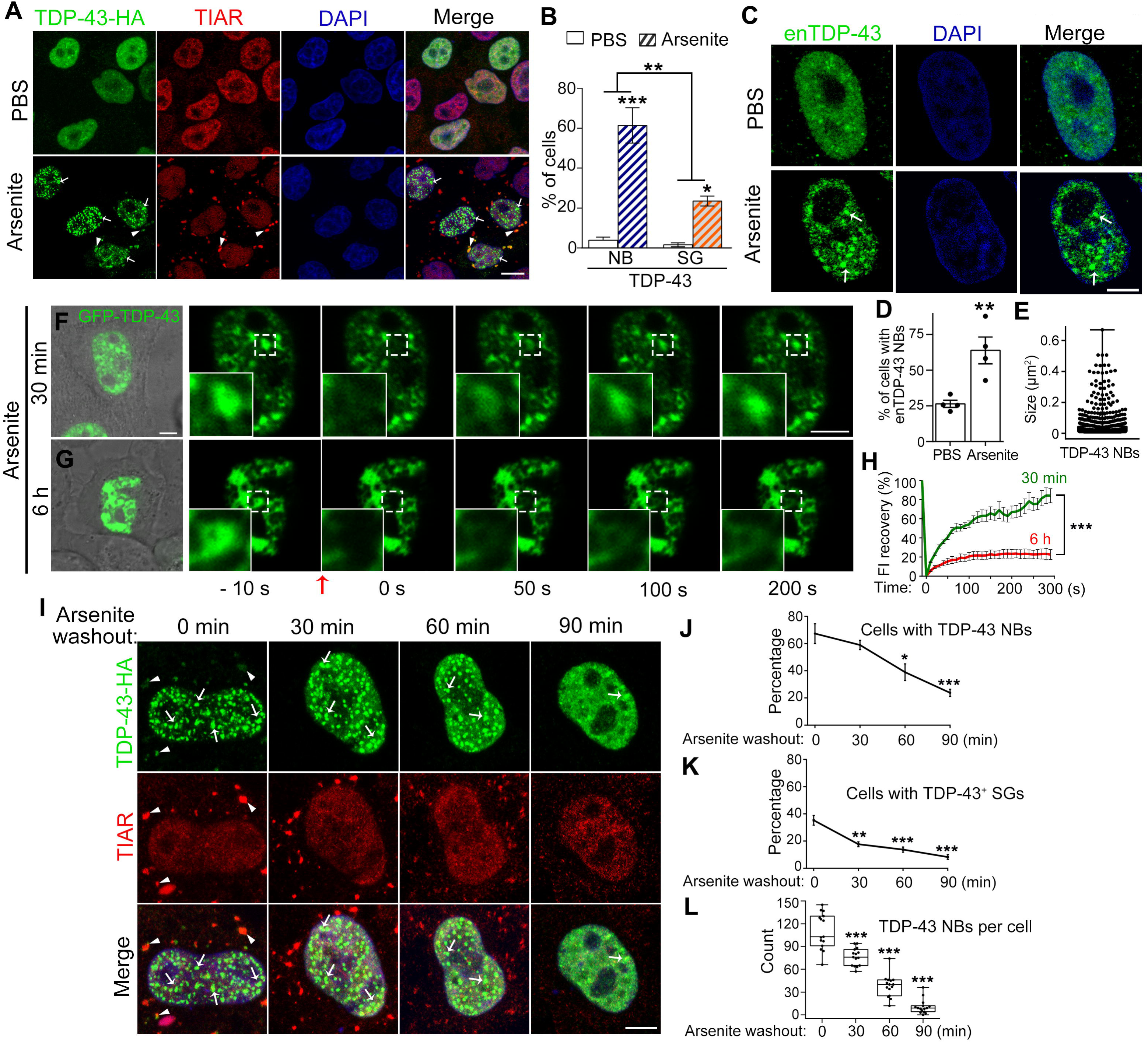
TDP-43 forms reversible, dynamic NBs in response to arsenic stress. **(A)** Representative confocal images of HeLa cells in the absence (PBS) or presence of arsenite (250 µM, 30 min) forming TDP-43 NBs (arrows) and TDP-43^+^ SGs (arrowheads) (also see Movie S1). All cells are transfected with TDP-43-HA (anti-HA) and stained for SGs with anti-TIAR; merged images with DAPI staining for DNA (blue) are shown. Arrows, TDP-43 NBs; arrowheads, SGs associated with TDP-43 (TDP-43^+^ SGs). **(B)** Quantification of the percentage of cells that show TDP-43 NBs or TDP-43^+^ SGs before and after arsenite the treatment in (A). **(C)** Representative images of the NBs formed by endogenous TDP-43 (enTDP-43) in HeLa cells upon arsenite treatment. **(D-E)** Quantifications of the percentage of cells in (C) showing stress-induced enTDP-43 NBs (D) and the size (µm^2^) of the NBs (E). **(F-G)** Representative images of the FRAP analysis of GFP-TDP-43 NBs in live cells after arsenite treatment for min (F) (also see Video S2) or 6 h (G). **(H)** The FRAP recovery curves of (F and G) are quantified by averaging the relative fluorescence intensity (FI) of TDP-43 NBs of similar size at indicated time. The relative FI of each TDP-43 NB prior to photobleaching is set to 100% and the time 0 refers to the time point right after photobleaching. **(I)** Representative images showing the disappearance of TDP-43 NBs and TDP-43^+^ SGs after arsenite washout. **(J-L)** Quantification of the percentage of cells showing TDP-43^+^ SGs (J), TDP-43 NBs (K) or the average count of TDP-43 NBs per cell (L) at indicated time points after arsenite washout in (I). Data are shown as mean ± SEM, except for (E) and (L) box-and-whisker plots with the value of each data measured. n = ~100 cells for each group in (B, D-E, J-K), n = 9 in (H), and n = ~15 cells for each group in (L), pooled results from 3 independent repeats. Statistic significance is determined at \**p* < 0.05, \*\**p* < 0.01, \*\*\**p* < 0.001 by Student’s t-test for comparison between PBS and arsenite within the same group and two-way ANOVA for comparison of the stress-induced changes between different groups in (B) Student’s t-test in (D), two-way ANOVA in (H), and one-way ANOVA in (J-L). Scale bars, 10 µm in (A) and 5 µm in (C, F, G and I).

A membraneless nuclear structure fulfills the requirements of NBs if it is: (1) microscopically visible, (2) enriched with specific nuclear factors, and (3) continuously exchanging the contents with the surrounding nucleoplasm (Stanek and Fox, 2017). Indeed, arsenite-induced TDP-43 nuclear granules were microscopically visible (Figures 1A-1E). Further examination by immunocytochemistry indicated that these arsenite-induced nuclear granules were colocalized or partially colocalized with several known types of NBs, especially paraspeckles labeled by SFPQ (Figure S1). Thus, the stress-induced TDP-43 nuclear granules were enriched of not only TDP-43 but also other nuclear factors such as paraspeckle proteins.

To examine if stress-induced TDP-43 nuclear granules were LD-like and could exchange with surrounding nucleoplasm, we performed the fluorescence recovery after photobleaching (FRAP) analysis *in live* cells using a green fluorescent protein (GFP)-tagged TDP-43 (GFP-TDP-43). To induce TDP-43 nuclear granules, cells were pre-treated with arsenite (250 µM) for 30 minutes before photobleaching. The fluorescence of GFP-TDP-43 nuclear granules was rapidly recovered after photobleaching, reaching ~50% of the pre-bleaching intensity in ~65 seconds (Figures 1F and 1H, and Movie S2), indicating that stress-induced TDP-43 nuclear granules are LD-like and highly dynamic. Together, they fulfill all the above three requirements and are thereafter called TDP-43 NBs in this study.

Next, we determined whether stress-induced TDP-43 NBs were reversible. In the arsenite washout assay, stress-induced TDP-43 NBs gradually disappeared and eventually recovered to nearly diffused pattern similar to that before the arsenite treatment (Figures 1I). Our quantitative analyses indicated that both the percentages of cells with TDP-43 NBs and the number of TDP-43 NBs per cell decreased in a time-dependent manner after arsenite washout (Figures 1J-1L). Thus, the arsenic stress-induced TDP-43 NBs are reversible. With prolonged stress such as the arsenite treatment (250 µM) for 6 h, the TDP-43 NBs were no longer dynamic and did not rapidly recover after photobleaching (Figures 1G and 1H).

### Formation of TDP-43 NBs as a general cellular stress mechanism

It was reported that TDP-43 formed reversible nuclear “aggregation” during heat shock (Udan-Johns et al., 2014), which together with the above arsenite-induced TDP-43 NBs raised the possibility that the assembly of TDP-43 NBs might be a general mechanism employed by the nucleus in response to stress. Consistent with this idea, we found that disturbing the nuclear homeostasis by inhibition of nuclear export with leptomycin B (LMB) also induced endogenous TDP-43 to form NBs (Figures 2A and 2B). Hence, as induced by arsenic stress (Figures 1C-1E), the formation of TDP-43 NBs was not an artificial effect simply due to TDP-43 overexpression.

**Figure 2.**
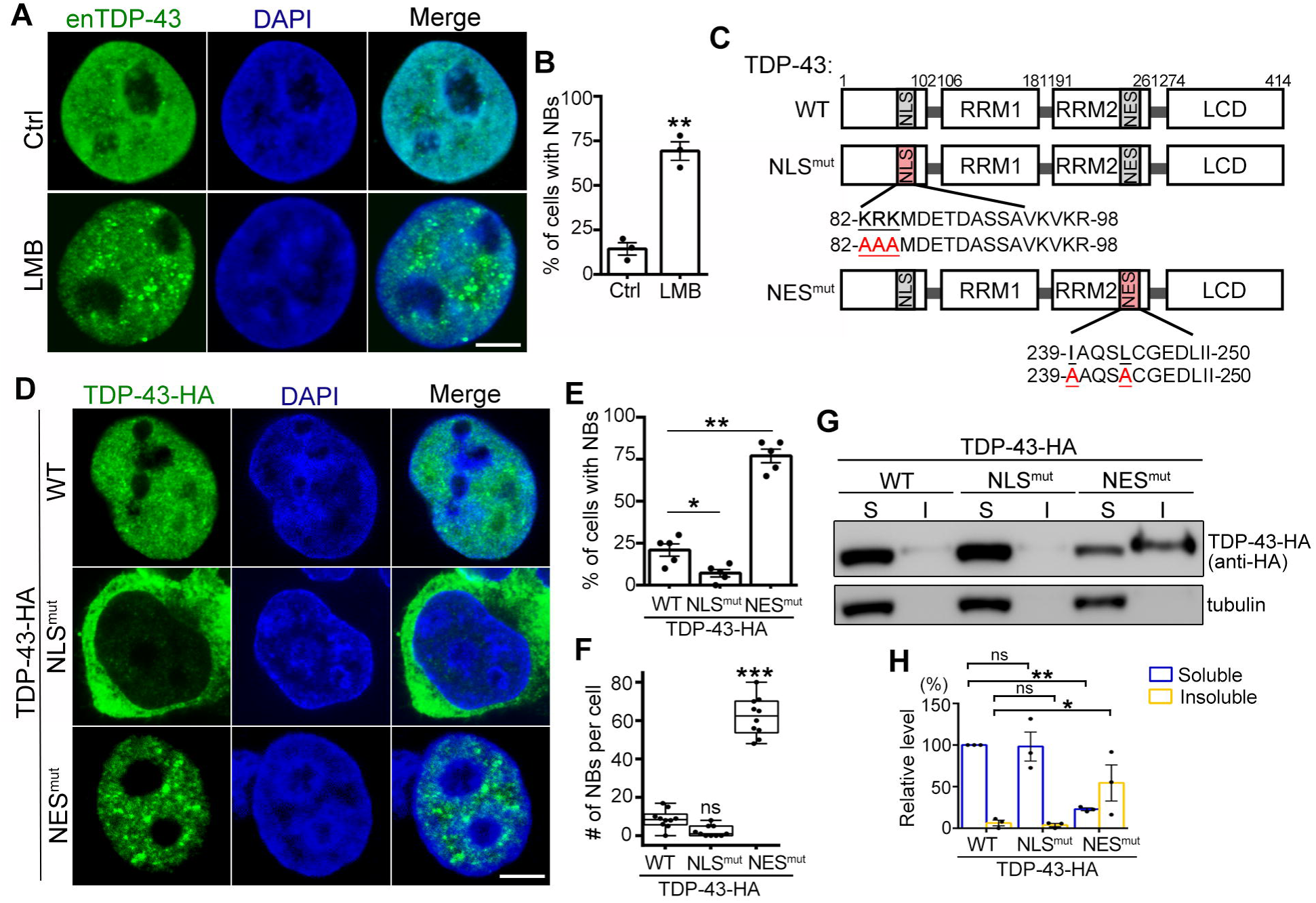
Inhibition of nuclear export induces the assembly of TDP-43 NBs. **(A-B)** Representative images of endogenous TDP-43 (enTDP-43) forming NBs upon LMB treatment (25 nM, 18 h). Percentage of cells showing distinct TDP-43 NBs in (A) is quantified in (B). **(C)** A diagram showing the major domains of human TDP-43 and the residues mutated in the localization signals. NLS, nuclear localization signal; RRM, RNA recognition motif; NES, nuclear export signal; LCD, low complexity domain. **(D)** Representative images of 293T cells transfected with WT, NLS^mut^ or NES^mut^ TDP-43. **(E-F)** The percentage of cells with NBs (E) and the average number of NBs per cell (F) are quantified. **(G-H)** Representative Western blot images (G) and quantifications (H) of WT, NLS^mut^ and NES^mut^ TDP-43 proteins in the soluble (S, supernatants in RIPA) and insoluble fractions (I, pellets resuspended in 9 M of urea). Mean ± SEM, except for (F) box-and-whisker plots. n = ~120 in (B, E) and ~10 cells in (F) for cells each condition or group from pooled results of 3 independent repeats, n = 3 in (G, H); \**p* < 0.05, \*\**p* < 0.01, \*\*\**p* < 0.001; ns, not significant; Student’s t-test in (B), one-way ANOVA in (E, F, H). Scale bar: 5 µm.

To determine whether blocking the nuclear export of TDP-43 itself was sufficient to induce the formation of TDP-43 NBs, we generated the NLS and NES mutants of TDP-43, namely NLS^mut^ and NES^mut^ (Figures 2C). Under normal conditions, wild-type (WT) TDP-43 was largely diffused in the nucleus with rare occurrence of spontaneous TDP-43 NBs (Figures 2D-2F). As expected, TDP-43-NLS^mut^ was predominantly cytoplasmic and thus the chance of the NLS^mut^ to form NBs was even lower than that of WT TDP-43 (Figures 2D and 2E). In contrast, the NES^mut^ was exclusively nuclear and formed TDP-43 NBs even in the absence of stress (Figures 2D-2F). Furthermore, the solubility of TDP-43-NES^mut^ protein was significantly reduced, as more than half of TDP-43-NES^mut^ protein was insoluble in radioimmunoprecipitation assay buffer (RIPA) (Figures 2G and 2H). Of note, the total protein levels (the sum of the soluble and insoluble fractions) of NLS^mut^ and NES^mut^ are not statistically different from that of WT TDP-43 (*p* = 0.8451 and 0.2835, respectively; one-way ANOVA).

The marked reduction in the solubility of the NES^mut^ protein raised the question whether the TDP-43-NES^mut^ NBs were still in a dynamic, LD-like state or had become solid aggregation. First, we performed the FRAP assay in *live* cells, which indicated that the NES^mut^ NBs were dynamic, as the fluorescence signal of GFP-TDP-43NES^mut^ rapidly recovered after photobleaching, just like the phenotype of the known nuclear body protein SFPQ (Figures S2A-S2B). Next, we examined the stability of WT and mutant TDP-43 by treating cells with protein synthesis inhibitor cycloheximide (CHX). We found that the RIPA-soluble TDP-43 protein was rather stable, as none of WT, NLS^mut^ or NES^mut^ TDP-43 in the soluble fractions showed significant turnover in 24 h after CHX treatment (Figures S2C-S2F). In contrast, the RIPA-insoluble fraction of TDP-43-NES^mut^ decreased rapidly upon CHX inhibition and became almost undetectable within 24 h (Figures S2C-S2F). Meanwhile, the CHX treatment led to disassembly of TDP-43-NES^mut^ NBs (Figures S2G and S2H). These results suggest that the RIPA-insoluble TDP-43-NES^mut^ NBs were dynamic and reversible, and therefore they were not “protein aggregates” as ones might presumed based on their punctate morphology and the detergent-insolubility. Of note, a recent study reported that stress promoted misfolded proteins to enter the phase-separated, LD-like nucleolus, which might function as a protein quality control mechanism in the nucleus (Frottin et al., 2019).

Next, we examined how disturbance of the cellular proteostasis impacted on TDP-43 NBs by inhibition of proteasome-mediated protein degradation with MG132. It led to a significant increase in the RIPA-insoluble fraction of NES^mut^ as well as WT TDP-43 protein (Figures S2I-S2L). In contrast, blocking autophagic flux with chloroquine (CQ) did not significantly affect the levels of insoluble TDP-43 (Figures S2M-S2P). Thus, unlike misfolded protein aggregates, the turnover of TDP-43 NBs did not rely on the autophagy-lysosomal pathway. Together, TDP-43 NBs are distinct from protein aggregates. Instead, they are dynamic, reversible membraneless subnuclear organelles that are sensitive and rapidly respond to changes in the microenvironment of the nucleoplasm.

### Formation of TDP-43 NBs alleviates the cytotoxicity in mammalian cells and fly neurons *in vivo*

We were keen to understand the functional significance of forming TDP-43 NBs in cells. Could it be a cellular “protective” mechanism like SGs? To answer this question, we examined how cells expressing WT, NLS^mut^ or NES^mut^ TDP-43 responded to cellular stress. Compared to WT TDP-43, cells transfected with the NLS^mut^ showed significantly increased cell death indicated by the propidium iodide (PI) staining, whereas the NES^mut^ was more resistant to arsenic stress (Figure S3A-S3D). Furthermore, we examined how the NB formation impacted on TDP-43 overexpression (OE)-mediated cytotoxicity *per* se. We found that OE of WT or NLS^mut^ but not NES^mut^ TDP-43 in human 293T cells significantly decreased the cell viability using the cell counting kit-8 (CCK-8) assay (Figures 3A) or by measuring the ATP levels (Figures 3B). These results were consistent with the previous studies that expression of cytoplasmic TDP-43 was more toxic than nuclear TDP-43 in flies (Miguel et al., 2011) and primary rat neurons (Barmada et al., 2010). However, the reduced cytotoxicity of nuclear TDP-43 had not been associated with the formation of NBs before.

**Figure 3.**
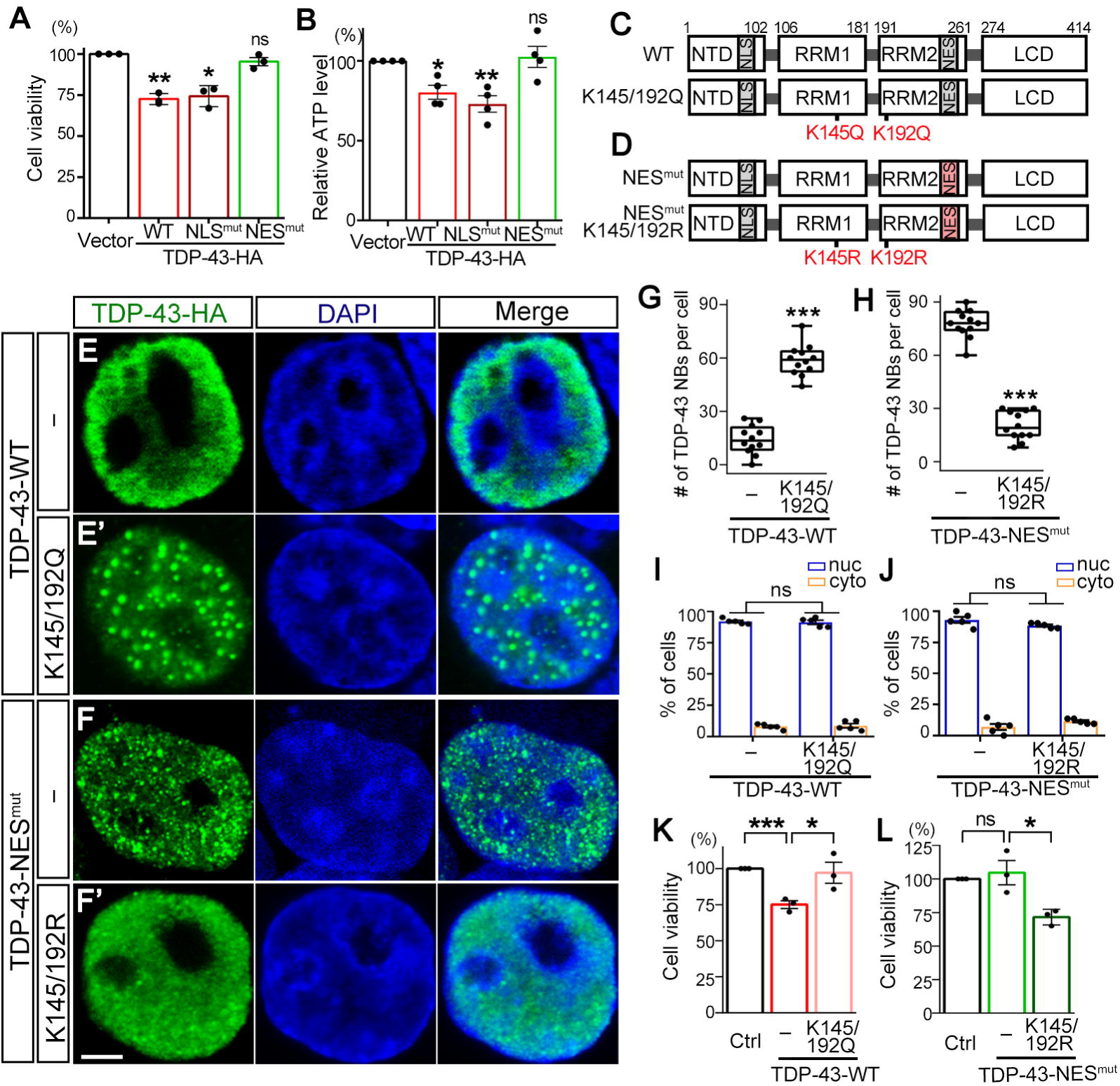
Formation of NBs mitigates TDP-43-mediated cytotoxicity. **(A)** OE of the WT or NLS^mut^ but not NES^mut^ TDP-43 in 293T cells decreases the cell viability, measured by a CCK-8 assay. **(B)** The relative ATP levels of the 293T cells transfected of WT, NLS^mut^ or NES^mut^ TDP-43 measured using a CellTiter-Glo® luminescent assay. **(C-D)** Diagrams showing the K145/192Q mutation in the WT TDP-43 and the K145/192R mutation in the NES^mut^ TDP-43, respectively. **(E-F’)** Representative confocal images of the WT (E) and “WT + K145/192Q” TDP-43 (E’), or the NES^mut^ (F) and “NES^mut^ + K145/192R” TDP-43 (F’) in 293T cells. **(G-H)** Quantifications of the average numbers of TDP-43 NBs per cell formed in each group in (E-F’) as indicated. **(I-J)** Percentages of cells with TDP-43 exclusively in the nucleus (nuc) or in both the nucleus and the cytoplasm (cyto) in (E-F’) are quantified. **(K-L)** The viability of cells expressing the WT or NES^mut^ TDP-43 with the indicated NB-promoting or NB-suppressing mutation is assessed using the CCK-8 assay. Mean ± SEM, except for (G-H) box-and-whisker plots; n = 3~4 in (A-B, K-L), n = ~12 cells in (G-H) and over 100 cells in (I-J) for each group from pooled results of 3 independent repeats; \**p* < 0.05, \*\**p* < 0.01, \*\*\**p* < 0.001; ns, not significant; one-way ANOVA in (A-B), Student’s t-test in (G-H, K-L), two-way ANOVA in (I-J). Scale bar: 5 µm.

To confirm whether TDP-43 form NBs *in vivo* and to examine how they impacted on neurodegeneration, we generated transgenic flies to express WT, NLS^mut^ or NES^mut^ human TDP-43 (hTDP-43) by ΦC31 integrase-mediated, site-specific integration (Groth et al., 2003; Bateman et al., 2006). The strains generated by this approach were different from the previously published NLS^mut^ and NES^mut^ hTDP-43 flies (Miguel et al., 2011), as in this study the different UAS-*hTDP*-43 transgenes were integrated into the fly genome at the same chromatin locus, which avoided the location effect on the expression levels of the transgenes among different strains and allowed direct comparison between the WT and the mutant TDP-43 flies. Immunohistochemistry analysis of the whole mount fly brains confirmed the predominant nuclear localization of the WT and NES^mut^ hTDP-43 in fly neurons (elav-Gal4), whereas the NLS^mut^ was largely cytoplasmic (Figure S4A). Consistent with the data in mammalian cells, the NES^mut^ hTDP-43 formed striking NBs in the nucleus of fly neurons (Figure S4A-S4B) and the solubility was significantly decreased (Figures S4C-S4D). We then examined the consequence of expressing WT, NLS^mut^ or NES^mut^ hTDP-43 in fly eyes to compare the degenerative morphological changes they caused (Figure S4E) and in motor neurons to compare the behavioral decline they induced (Figure S4F). In both experiments, WT TDP-43 flies exhibited marked age-dependent degenerative phenotypes and the NLS^mut^ flies were significantly worse. In contrast, the neurotoxicity of TDP-43-NES^mut^ was substantially milder, especially in the ALS-related functional assay, the climbing capability of the NES^mut^ flies was not statistically different from that of the control flies (UAS-Luciferase) at all time points examined (Figure S4F).

To further confirm that the formation of TDP-43 NBs was indeed required for the cytotoxicity-antagonizing effect rather than simply restricting TDP-43 in the nucleus, we sought for alternative approaches to induce TDP-43 NBs without disrupting its nucleocytoplasmic transport. Acetylation of TDP-43 at K145 and K192 was previously shown to regulate RNA binding and affect TDP-43 aggregation (Cohen et al., 2015). The acetylation-mimic mutation (K145/192Q) induced whereas the acetylation-deficient mutation (K145/192R) suppressed the formation of TDP-43 protein inclusions (Cohen et al., 2015). Hence, we generated K145/192Q mutation in WT TDP-43 (Figures 3C), which led to the assembly of TDP-43 NBs in the absence of cellular stress (Figures 3E-3E’ and 3G). On the other hand, we introduced K145/192R mutation into TDP-43-NES^mut^ (Figures 3D), which abolished the spontaneous TDP-43-NES^mut^ NBs (Figures 3F-3F’ and 3H). In addition, we confirmed that neither K145/192Q nor K145/192R significantly altered the subcellular distribution of TDP-43 (Figures 3I and 3J). Consistent with our hypothesis, the NB-forming K145/192Q mutation alleviated the cytotoxicity of WT TDP-43 (Figures 3K), whereas K145/192R abolished the NB-forming capability and made the originally non-toxic TDP-43-NES^mut^ to manifest marked cytotoxicity (Figures 3L). Thus, the formation of TDP-43 NBs can mitigate cytotoxicity, which might help cells to survive stressed or diseased conditions.

### The role of the major functional domains of TDP-43 in the assembly of NBs

We went on to investigate how TDP-43 NBs were assembled. First, to understand the function of each major domain in TDP-43 NB formation, we generated truncated TDP-43, namely ΔLCD, ΔRRM1 and ΔRRM2 (Figure S5A). Both WT and ΔLCD TDP-43 proteins were mostly nuclear and soluble (Figures S5B and S5C); however, ΔLCD did not form NBs in response to stress (Figures S5F to S5G’), confirming a major role of the LCD domain in promoting the assembly of RNP granules and protein aggregation (Molliex et al., 2015; Conicella et al., 2016). Next, we examined the impact of RRM1 or RRM2 on the assembly of TDP-43 NBs. TDP-43-ΔRRM1 was soluble (Figure S5D), whereas ΔRRM2 showed a remarkable increase of insolubility (Figure S5E). More interestingly, TDP-43-ΔRRM1 formed large, ring-shaped structures in the nucleus without any stress (Figure S5H-S5H’); while ΔRRM2 looked similar to WT TDP-43 without stress and showed a grainy appearance with LMB treatment (Figures S5I-S5I’).

To further characterize the nuclear structures formed by WT, ΔRRM1 and ΔRRM2 TDP-43, we utilized the Leica LIGHTNING SP8 confocal microscope to capture multicolor images in super-resolution down to 120 nm. We found that WT TDP-43 formed solid, mid-size NBs; ΔRRM1 was much larger, ring-shaped; and ΔRRM2 displayed a mesh-like structure with numerous smaller NBs (Figures 4A and 4B). Super-resolution three-dimensional (3D) rendering revealed that WT and ΔRRM2 TDP-43 NBs were mostly in oval or sometimes cylinder shapes, while ΔRRM1 formed large, hollow “pillars”. Further measurement of the diameters confirmed that ΔRRM1 “rings” was drastically larger whereas ΔRRM2 NBs was slightly smaller than WT TDP-43 NBs (Figures 4C).

**Figure 4.**
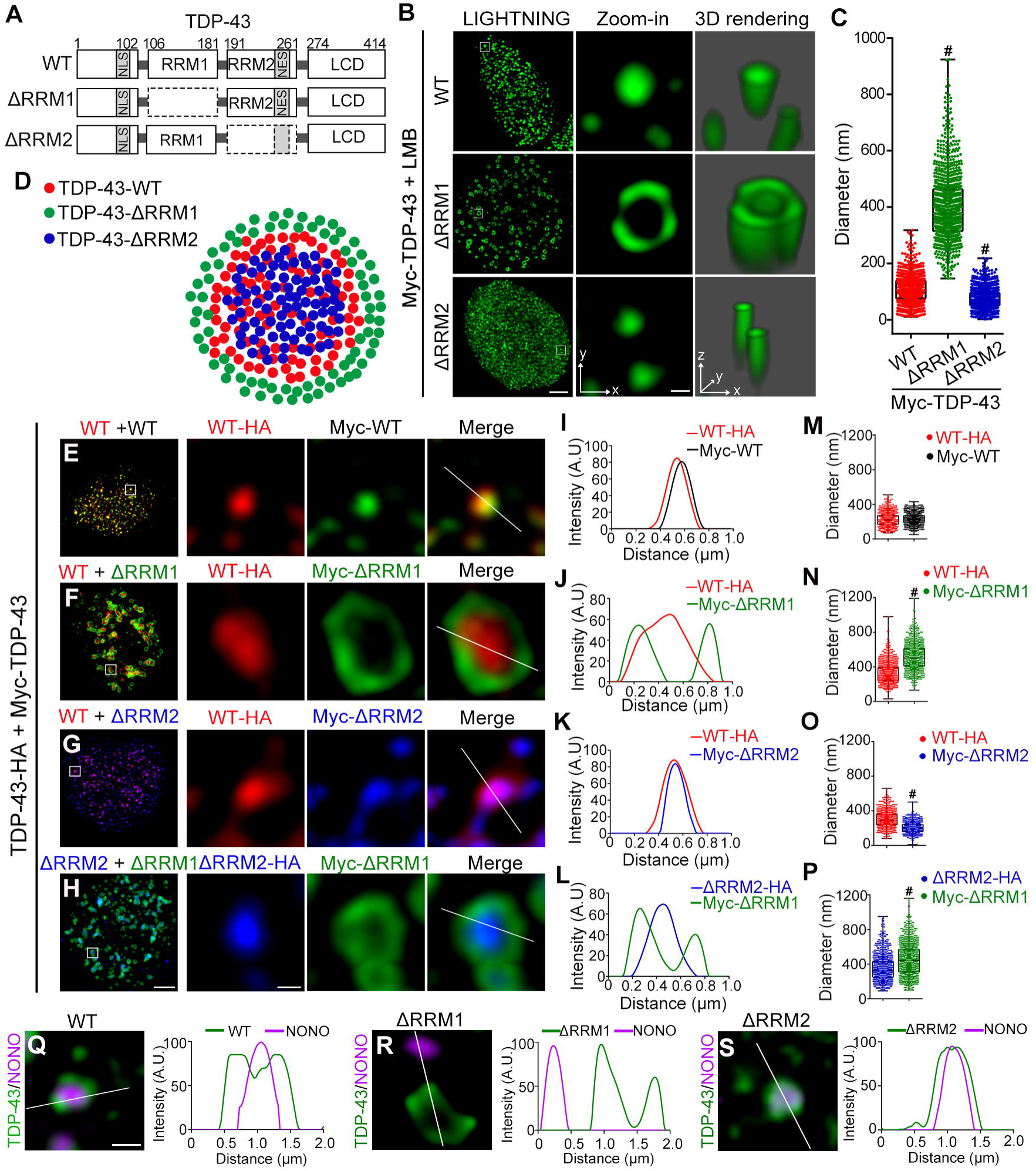
The RRM1 and RRM2 function complementarily to maintain the core-shell organization of TDP-43 NBs. **(A)** A diagram showing the major functional domains of WT TDP-43 and the ΔRRM1 and ΔRRM2 mutants. **(B)** LIGHTNING super-resolution microscopy of 293T cells transfected with Myc-TDP-43-WT, ΔRRM1 or ΔRRM2 and treated with LMB. Higher magnification images (zoom-ins) and 3D renderings of the boxed areas are shown. **(C)** The average diameters of the WT, ΔRRM1 and ΔRRM2 TDP-43 nuclear structures in (B) are quantified. **(D)** A schematic model of the core-shell organization of TDP-43 NBs (red) and the distinct distribution of ΔRRM1 (green) and ΔRRM2 (blue). **(E-H)** LIGHTNING multicolor super-resolution microscopy and higher magnification images of representative LMB-induced TDP-43 NBs. Cells are co-transfected with a combination of HA-or Myc-tagged WT, ΔRRM1 or ΔRRM2 TDP-43 as indicated. **(I-P)** The intensity profiles (I-L) along the indicated lines and the average diameters (M-P) of TDP-43 NBs in each group are shown. **(Q-S)** Representative super-resolution microscopy images and the intensity line analyses of WT (Q), ΔRRM1 (R) or ΔRRM2 (S) TDP-43 nuclear structures (green) co-immunostained with the paraspeckle protein NONO (purple) are shown. The box-and-whisker plots with the value of each NB measured are shown. n = ~900 NBs in (C, M-P) for each group from pooled results of at least 3 independent repeats; #p < 0.0001; one-way ANOVA in (C) and Student’s t-test in (M-P). Scale bars: 5 µm in (B), 2 µm in (E-H), 200 nm in the zoom-ins, and nm in (Q-S).

Given the distinct morphology of the nuclear structures formed by ΔRRM1 and ΔRRM2 TDP-43, it was tempting to propose that TDP-43 NBs, similar to paraspeckles (West et al., 2016), might have a “core-shell” organization (Figures 4D). Moreover, we hypothesized that the two RRMs might function antagonistically to keep the shape and the size of TDP-43 NBs. In other words, the tension maintained by the two RRMs might prevent TDP-43 NBs from expanding unrestrainedly to become huge nuclear bubbles as well as from condensing all into the core to become insoluble aggregates. To test this hypothesis, we co-expressed HA-tagged WT TDP-43 with Myc-tagged WT, ΔRRM1 or ΔRRM2 TDP-43 and induced the NB formation in 293T cells by LMB (Figures 4E-4H). The super-resolution images showed that ΔRRM1 formed a ring-shaped shell wrapping the co-expressed WT TDP-43 NBs (Figures 4F and 4J) and the size of the NBs was much larger (Figures 4N). The increased diameter was not simply due to overexpression of two folds of TDP-43 protein, as co-expression of WT TDP-43-HA and WT Myc-TDP-43 did not manifest such effect (Figures 4E, 4I, and 4M). Consistent with our hypothesis, ΔRRM2 co-expressed with WT TDP-43 was mainly located in the core of the stress-induced NBs (Figures 4G and 4K), and the size was significantly smaller than WT TDP-43 NBs (Figures 4O). The core-shell organization could also be clearly seen when ΔRRM1 and ΔRRM2 TDP-43 were co-expressed (Figures 4H, 4L, and 4P).

Next, we characterized the co-localization profiles of the above TDP-43 nuclear structures with paraspeckles by co-immunostaining of NONO and super-resolution microscopy. Of note, NONO is a paraspeckle protein known to reside in the core of paraspeckles (West et al., 2016). Consistently, NONO also showed up in the core of WT TDP-43 NBs (Figures 4Q), which was more obvious when overlaid with ΔRRM2 (Figures 4S). Interestingly, although the ring-shaped structure of ΔRRM1 co-localized and surrounded WT TDP-43 NBs (Figures 4F and 4J), it no longer co-localized with paraspeckles labeled by NONO (Figures 4R). These results indicate that TDP-43 requires the RRM1 domain to be associated with paraspeckles. Also, they confirm that TDP-43 NBs are not homogenous, although a significant subpopulation overlapped with paraspeckles (Figures S1). Together, these data suggest that RRM1 and RRM2 play distinct functions in the assembly and maintenance of TDP-43 NBs, possibly by interacting with different RNAs and/or different RBPs.

### RNA suppresses TDP-43 *in vitro* de-mixing via the RRMs

The dynamic, reversible and membraneless characteristics of TDP-43 NBs suggested that LLPS might be involved in the formation of TDP-43 NBs. Indeed, purified full-length (FL) TDP-43 protein formed LDs *in vitro* in a dose-dependent manner and the LCD played a major role in promoting TDP-43 LLPS (Figures S6A-S6D). Nevertheless, *in vitro* de-mixing of the C-terminal LCD truncation (TDP-43^1-274^) was evident at a higher protein concentration (≥50 µM) (Figure S6D), which was consistent with the previous report that the N-terminal domains of TDP-43 could drive LLPS *in vivo* (Schmidt and Rohatgi, 2016). Next, we examined how RNA impacted on TDP-43 LLPS by adding total RNAs extracted from HeLa cells into the *in vitro* de-mixing system (Figures S6E and S6F). Total RNAs (500 ng/µl) markedly reduced the LLPS of FL TDP-43 (Figure S6E), whereas much lower concentrations of total RNAs (≥100 ng/µl) were sufficient to suppress the LLPS of TDP-43^1-274^ (Figure S6F). Of note, the presence of the LCD made the purified FL TDP-43 protein extremely insoluble, for which a sumo tag was needed all the time to keep FL TDP-43 protein from precipitation *in vitro*. However, the solubilizing sumo tag made it difficult to assess and compare the role of the two RRMs in mediating RNA suppression, as large SUMO-TDP-43 droplets were tricky to form or maintain (Figures S6G-S6I). Thus, we used the TDP-43^1-274^ protein (which did not require the sumo tag to stay soluble) in the subsequent experiments (Figures S6J-S6L and Figure 5). In addition, we confirmed that the RNA-binding affinity was significantly reduced in ΔRRM1 and ΔRRM2 TDP-43^1-274^ (Figures S6M-S6O).

Both WT TDP-43^1-274^ and the two RRM mutants formed LDs in a dose-dependent manner in the *in vitro* LLPS system, though the de-mixing of ΔRRM1 TDP-43^1-274^ was less robust (Figures S6J-S6L). To facilitate the subsequent evaluation, we lowered the NaCl concentration and increased the crowding agent PEG, which allowed larger TDP-43 LDs to form at the beginning of the suppression assay (Figures 5B-5D). We found that total RNAs at a concentration of 100 ng/µl markedly reduced the size and number of WT TDP-43^1-274^ LDs. ΔRRM1 TDP-43^1-274^ (50 µM) only formed small LDs, which initially made us presume the LLPS of ΔRRM1 would be more easily suppressed by total RNAs. To our surprise, a much higher concentration of total RNAs (≥500 ng/µl) was required (Figures 5C). We also tested ΔRRM1 at a higher concentration (100 µM) to allow large ΔRRM1 LDs to form at the beginning, and the results confirmed the drastically reduced RNA suppression of the LLPS of ΔRRM1 TDP-43^1-274^ (total RNAs ≥250 ng/µl) (Figure 5C’). The RNA suppression of the LLPS of ΔRRM2 TDP-43^1-274^ was also reduced, but to a less extent than that of ΔRRM1 (100~250 ng/µl total RNAs) (Figures 5B-5D). It was worth noting that ΔRRM1 TDP-43^1-274^ showed both decreased LLPS capacity and reduced sensitivity to RNA suppression, indicating that a RBP with lower tendency to phase separate does not necessarily mean that RNA would more potently suppress its LLPS. Rather, the regulation by RNAs is likely to be specific and different depending on the type and structure of the RNAs.

**Figure 5.**
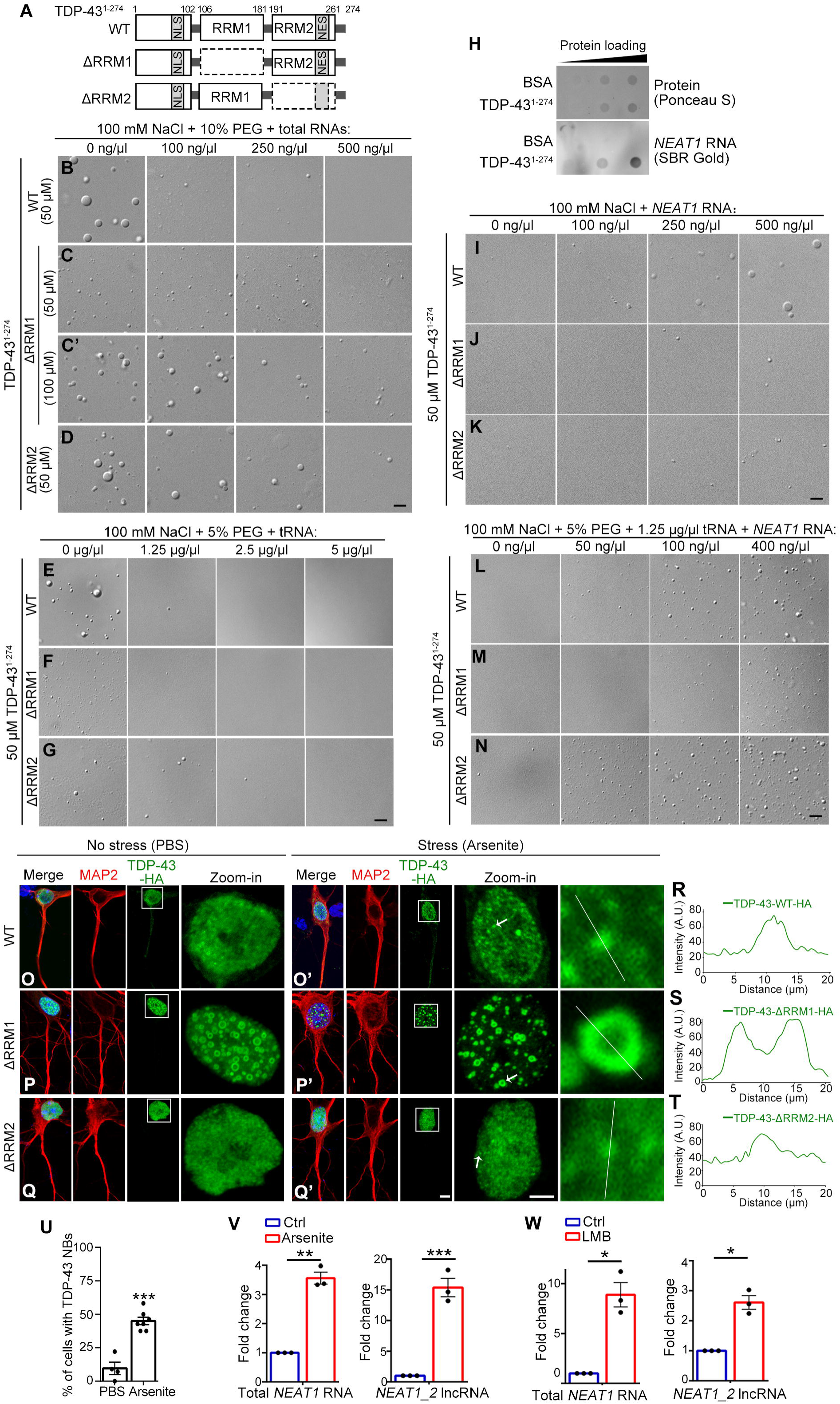
*NEAT1* RNA promotes TDP-43 LLPS *in vitro* and is upregulated in stressed neurons. **(A)** A diagram showing the WT, ΔRRM1 and ΔRRM2 of TDP-43^1-274^. **(B-K)** The total RNA extracts (B-D), tRNA (E-G), or lncRNA *NEAT1* (I-K) are added into the *in vitro* LLPS assay of WT, ΔRRM1 and ΔRRM2 TDP-431-273 as indicated. C’, a higher concentration of ΔRRM1 is also included in order to start the suppression assay with large LDs as those of WT TDP-43. H, representative images of the *in vitro* dot-blot assay confirming direct binding of *NEAT1* RNA to TDP-43 protein. The bound RNA is visualized using the SYBR® Gold Nucleic Acid Gel Stain kit and the protein loaded is stained by Ponceau S. Bovine serum albumin (BSA) is used as a negative binding control. **(L-N)** tRNA-mediated suppression of TDP-43 LLPS can be antagonized by increasing the level of *NEAT1* RNA. The concentrations of NaCl, the crowding agent PEG, TDP-43 proteins and RNAs used in the *in vitro* assays are indicated. **(O-Q’)** Representative images of primary mouse cortical neurons infected with WT (O-O’), ΔRRM1 (P-P’) and ΔRRM2 (Q-Q’) TDP-43 treated with PBS or arsenite as indicated. TDP-43-HA (green), microtubule associated protein 2 (MAP2) to mark neuronal perikarya and dendrites (red), and merged images with DAPI staining to label the nucleus (blue). Zoom-in images of the boxed areas are shown. **(R-T)** Higher magnification images and the intensity line analyses of the nuclear structures pointed by the arrows in O’-Q’ are shown. **(U)** The percentage of primary mouse neurons forming WT TDP-43 NBs in response to arsenic stress in (O-O’) is quantified. **(V-W)** Cellular stress induced by arsenite (V) or LMB (W) increases the levels of both total *NEAT1* RNA and the *NEAT1_2* isoform in mouse primary neurons compared to the vehicle control. Mean ± SEM; n = ~100 cells each group from pooled results of 3 independent repeats in (U), n = 3 in (V-W); \**p* < 0.05, \*\**p* < 0.01, \*\*\**p* < 0.001; Student’s t-test. Scale bars: 2 µm in (B-G, I-N), 5 µm in (O-Q’) and 2 µm in the zoom-ins.

tRNA reduced the LLPS of FUS *in vitro* (Maharana et al., 2018), which also potently suppressed the *in vitro* de-mixing of TDP-43 (Figures 5E). Further, we found that WT and ΔRRM1 TDP-43^1-274^ showed similar sensitivity to tRNA (≥1.25 µg/µl) (Figures 5E and 5F), whereas ΔRRM2 required a higher concentration of tRNA (≥2.5 µg/µl) to reach the similar extent of suppression (Figures 5G). It should be pointed out that, unlike the response to total RNAs, ΔRRM1 TDP-43^1-274^ did not exhibit a greatly decreased sensitivity to tRNA suppression (Figures 5C and 5F). Thus, the lack of a strong suppression of the LLPS of ΔRRM1 by total RNAs was not simply because suppression of small LDs was difficult to manifest or detect. Taken together, RRM1 appeared to play a major role in mediating total RNA suppression of TDP-43 LLPS (Figures 5B-5D), whereas RRM2 might be specifically involved in the suppression by certain RNAs such as tRNA (Figures 5E-5G).

### Paraspeckle RNA *NEAT1* promotes TDP-43 LLPS and is upregulated in stressed neurons

Recent works have revealed both positive and negative regulation of LLPS of prion-like RBPs by RNAs (Guo and Shorter, 2015). Since stress-induced TDP-43 NBs were partially co-localized with paraspeckles (Figures S1A and S1B), we examined whether the paraspeckle scaffolding RNA *NEAT1* impacted on the phase behavior of TDP-43. We showed that *NEAT1* RNA could directly bind to TDP-43 protein in the *in vitro* dot-blot assay (Figures 5H). Further, addition of *NEAT1* RNA into the *in vitro* de-mixing system promoted TDP-43 LLPS in a dose-dependent manner (Figures 5I), and this effect involved both RRM1 and RRM2 (Figures 5J and 5K). Furthermore, we revealed that increasing concentrations of *NEAT1* antagonized the suppressive environment generated by tRNA in the *in vitro* LLPS assay (Figures 5L). Moreover, the promotion of TDP-43 LLPS by *NEAT1* was markedly reduced in ΔRRM1 but enhanced in ΔRRM2 (Figures 5M and 5N), suggesting that, in a complex system containing both positive and negative regulatory RNAs, such as in the nucleoplasm, the RRM1 of TDP-43 may play a major role in mediating the *NEAT1*-promotion of TDP-43 LLPS whereas the RRM2 may be more involved in the tRNA-mediated suppression.

Furthermore, we examined the RNA-binding affinity of the NES^mut^ and found that it was significantly lower than that of WT TDP-43 (Figures S3G-S3H), which confirmed the previous reports that the NES^mut^ impaired RNA binding of TDP-43 (Lukavsky et al., 2013; Flores et al., 2019). Consistently, the K145/192Q mutation also showed reduced RNA binding (Figures S3E-S3F), and the suppression of TDP-43 LLPS by total RNAs was decreased with the K145/192Q (Figures S3I-S3J). Interestingly, the K145/192R mutation did not appear to improve RNA binding (Figure S3E-S3H) or enhance the suppression of TDP-43 LLPS by total RNAs (Figure S3K). Instead, it disrupted *NEAT1*-mediated nucleation of TDP-43 droplets (Figures S3L-S3N), which might “complement” the lack of RNA suppression in the NES^mut^ and thus reversed the spontaneous NB formation in TDP-43-NES^mut^-K145/192R (Figures 3F-3F’ and 3H) and regained cytotoxicity (Figures 3L). These data suggest a vital role of *NEAT1* in the formation of TDP-43 NBs.

Next, we extended our investigation of TDP-43 NBs and *NEAT1* to mammalian neurons. We confirmed that TDP-43 formed NBs in mouse primary neurons in response to stress (Figures 5O-5O’, 5R and 5U). Consistent with the phenotypes in HeLa and 293T cells, ΔRRM1 showed large ring-shaped structures in both non-stressed and stressed mouse neurons (Figures 5P-5P’ and 5S), whereas ΔRRM2 displayed a grainy appearance when neurons were treated with arsenite (Figures 5Q-5Q’ and 5T). More importantly, we found that the levels of both total *NEAT1* RNA and the lncRNA isoform *NEAT1_2* were dramatically increased in mouse primary neurons treated with arsenite (Figures 5V) as well as LMB (Figures 5W). Together with the *in vitro* LLPS data (Figures 5L-5N), these data suggest that stress may trigger an upregulation of *NEAT1* RNA, which provides nucleation scaffolds to condense TDP-43 and possibly other NB components by promoting the LLPS, thereby antagonizing the suppressive environment of the nucleoplasm and causing the formation of TDP-43 NBs.

### ALS-associated D169G mutation of TDP-43 shows a specific defect in NB formation

This study was launched by the discovery that TDP-43 formed dynamic and reversible NBs in stressed cells and the formation of NBs alleviated TDP-43-mediated cytotoxicity. To further investigate the disease relevance of TDP-43 NBs, we examined several known ALS-causing mutations in TDP-43. Interestingly, we found that the mutation of D169G (640AT) within the RRM1 (Kabashi et al., 2008; and Figures 6A-6B) drastically reduced the assembly of TDP-43 NBs induced by either arsenite (250 µM, min) (Figures 6C-6E) or LMB (25 nM, 12 h) (Figures S7A-S7C). In contrast, the assembly of SGs or the recruitment of TDP-43-D169G to SGs was not diminished at this condition (Figures 6C and 6F). Unlike D169G, the disease-associated mutations in the LCD region such as Q331K and M337V did not hindered the stress-induced assembly of TDP-43 NBs (Figures S7D-S7G). Thus, the D169G mutation specifically impaired the assembly of stress-induced TDP-43 NBs. In addition, the difference between D169G and other disease mutations enriched in the LCD also suggests that although TDP-43 NBs and SGs are both phase-separated, membraneless RNP granules, the exact mechanisms regulating their formation can be different.

**Figure 6.**
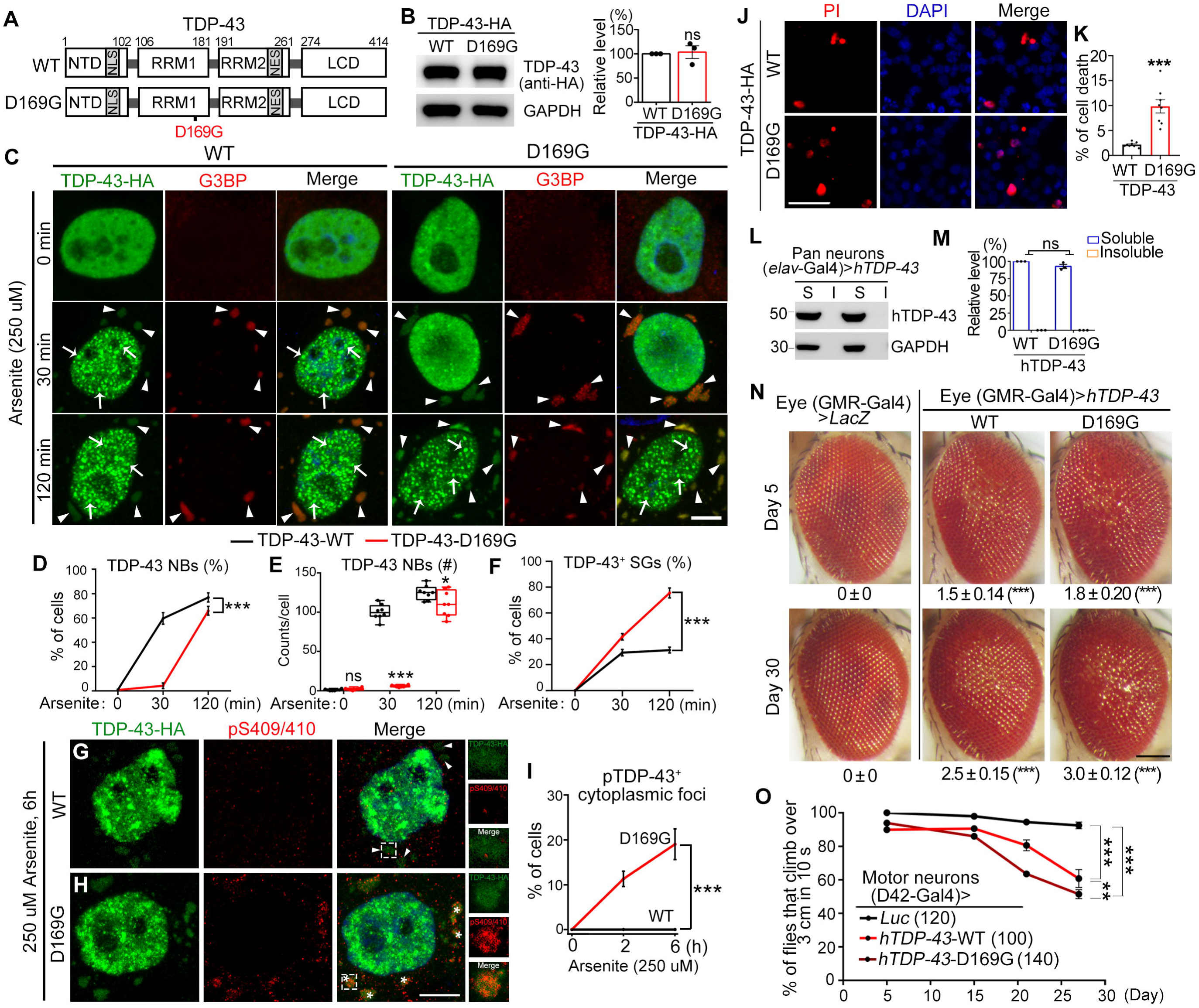
TDP-43-D169G impairs NB assembly, accumulates pathological cytoplasmic foci and potentiates the cytotoxicity of TDP-43 in prolonged stress and during aging. **(A)** A diagram showing WT and D169G mutant TDP-43. **(B)** Representative Western blot image and quantification confirming the expression levels of WT and D169G TDP-43 in HeLa cells. **(C)** Representative confocal images of HeLa cells transfected with WT or D169G TDP-43, treated with arsenite for 0 min (PBS), min or 120 min as indicated. Green, TDP-43-HA (anti-HA); Red, anti-G3BP for SGs; Blue, DAPI staining for DNA (nucleus). **(D-F)** The percentage of cells with TDP-43 NBs (D), the average count of TDP-43 NBs per cell (E) and the percentage of cells with TDP-43^+^ SGs (F) at indicated time after arsenite treatment in (C). **(G-H)** Confocal images of representative single cells expressing WT (G) or D169G TDP-43-HA (H) that form cytoplasmic foci (green) after arsenite treatment for 6 h were co-stained for pS409/410 (red) and DAPI for DNA (blue). Higher magnification images of the boxed areas are shown on the side. **(I)** The percentage of cells with pS409/10+ TDP-43 cytoplasmic foci at indicated times during prolonged stress (arsenite, 250 µM) is quantified. Arrows, TDP-43 NBs; arrowheads, TDP-43^+^ SGs; asterisks, pTDP-43^+^ cytoplasmic foci. **(J-K)** Representative images (J) and quantification (K) of PI staining to evaluate the death of cells expressing WT or D169G TDP-43. **(L-M)** Representative images (L) and quantification (M) of the Western blot analysis confirming the expression levels and solubility of hTDP-43-D169G in the transgenic flies. **(N)** Representative z-stack images of the fly eye (driven by a GMR-Gal4 driver) expressing WT or D169G hTDP-43 at indicated time points. The degeneration severity is assessed by the rough surface, swelling and loss of pigment cells of the compound eyes. The average degeneration score (mean ± SEM) and the statistic significance (compared to the UAS-lacZ control flies) are indicated at the bottom of each group. **(O)** The climbing capability of the flies expressing WT or D169G hTDP-43 in motor neurons (D42-Gal4) is evaluated as percentage of flies climbing over 3 cm in 10 seconds. The UAS-luciferase fly line is used as a control. The numbers of flies tested in each group are indicated. Mean ± SEM, except for (E) box-and-whisker plots; n = 3 in (B, M), n = ~100 cells in (D, F, I), ~8 cells in (E) and ~2000 cells in (K) for each group from pooled results of 3 independent repeats, n = 13~16 eyes each group in (N); \**p* < 0.05, \*\**p* < 0.01, \*\*\**p* < 0.001; ns, not significant; Student’s t-test in (B, E, K, M), one-way ANOVA in (N) and two-way ANOVA in (D, F, I, O). Scale bars: 5 µm in (C, G-H), µm in (J) and 100 µm in (N).

Next, we increased the duration of arsenite treatment (250 µM, 120 min) to test if D169G could eventually form TDP-43 NBs with prolonged stress, which indeed occurred (Figures 6C-6E). However, what was unexpected and surprised us was that there were staggeringly more cells forming TDP-43^+^ SGs in D169G (75.6% ± 3.8%) than WT TDP-43 (31.3 ± 2.3%) with prolonged stress (Figures 6C and 6F). It has been proposed that TDP-43 recruited to SGs becomes phosphorylated, which promotes aberrant phase transition of SGs to cytoplasmic protein inclusions (Li et al., 2013). And, phosphorylation of TDP-43 (pTDP-43) at S409/410 (pS409/410) is a disease-specific marker of TDP-43 aggregates (Hasegawa et al., 2008; Neumann et al., 2009). Immunostaining for pS409/410 revealed that the NB-forming defective D169G formed pTDP-43 cytoplasmic foci in prolonged stress (arsenite 250 µM, 6 h), which was not observed in WT TDP-43 under the same condition (Figures 6G-6I). Of note, like WT TDP-43 (Figures 6G), D169G in the nucleus was absent of pS409/410 staining even with prolonged stress (Figures 6H), indicating that TDP-43-D169G was not generally hyperphosphorylated but rather only when it formed abnormal cytoplasmic foci. Moreover, cells expressing D169G showed significantly more death during prolonged stress (arsenite 100 µM, 12 h) than cells expressing WT TDP-43 (Figures 6J-6K). These results suggest that the assembly of TDP-43 NBs acts as an important stress response mechanism, which may lower the demand for the formation of SGs and prevent excessive cytoplasmic translocation of TDP-43, therefore reducing the chance of aberrant phase transition and the accumulation of pathogenic pTDP-43 inclusions in the cytoplasm.

To further assess the cytotoxicity of the D169G mutation in an *in vivo* setting, we generated transgenic flies expressing *hTDP-43-D169G* (Figures 6L-6M). These flies exhibited age-dependent degeneration of the eyes (Figures 6N), and the enhanced neurotoxicity was more evident when the D169G mutant was expressed in fly motor neurons – the climbing capability showed an accelerated decline compared to the flies expressing *WT-hTDP-43*, especially at later time points in aged flies (Figures 6O, Day 21 and Day 27). As aging is associated with cumulative stress and is a risk factor for neurodegenerative diseases (Lopez-Otin et al., 2013; Hou et al., 2019), these results further support the idea that the D169G mutation makes cells more prone to stress, which may underlie its pathogenesis in the ALS disease.

### The D169G mutation diminishes lncRNA *NEAT1*-mediated TDP-43 LLPS *in vitro* and NB formation in cells

To understand the molecular basis for the NB-forming deficit of D169G, we then examined the phase separation behavior of TDP-43-D169G. The results of the *in vitro* LLPS assay showed that D169G TDP-43^1-274^ protein formed LDs in a dose-dependent manner just like that of WT (Figures 7A-7B), which indicated that the ability of D169G to phase separate was not markedly changed. The dot-blot assay confirmed the previous report (Kuo et al., 2014) the binding affinity of D169G to total RNAs was not reduced compared to that of WT TDP-43 (Figures S7H-S7J). And, suppression of TDP-43 LLPS by total RNA extracts showed no marked difference in D169G compared to that of WT TDP-43^1-274^ (Figures 7C and 7D). Thus, the general regulation of TDP-43 RNP granules by total RNAs remained largely intact in D169G, which was consistent with the fact that D169G did not hinder SG formation in stressed cells (Figures 6C and 6F).

**Figure 7.**
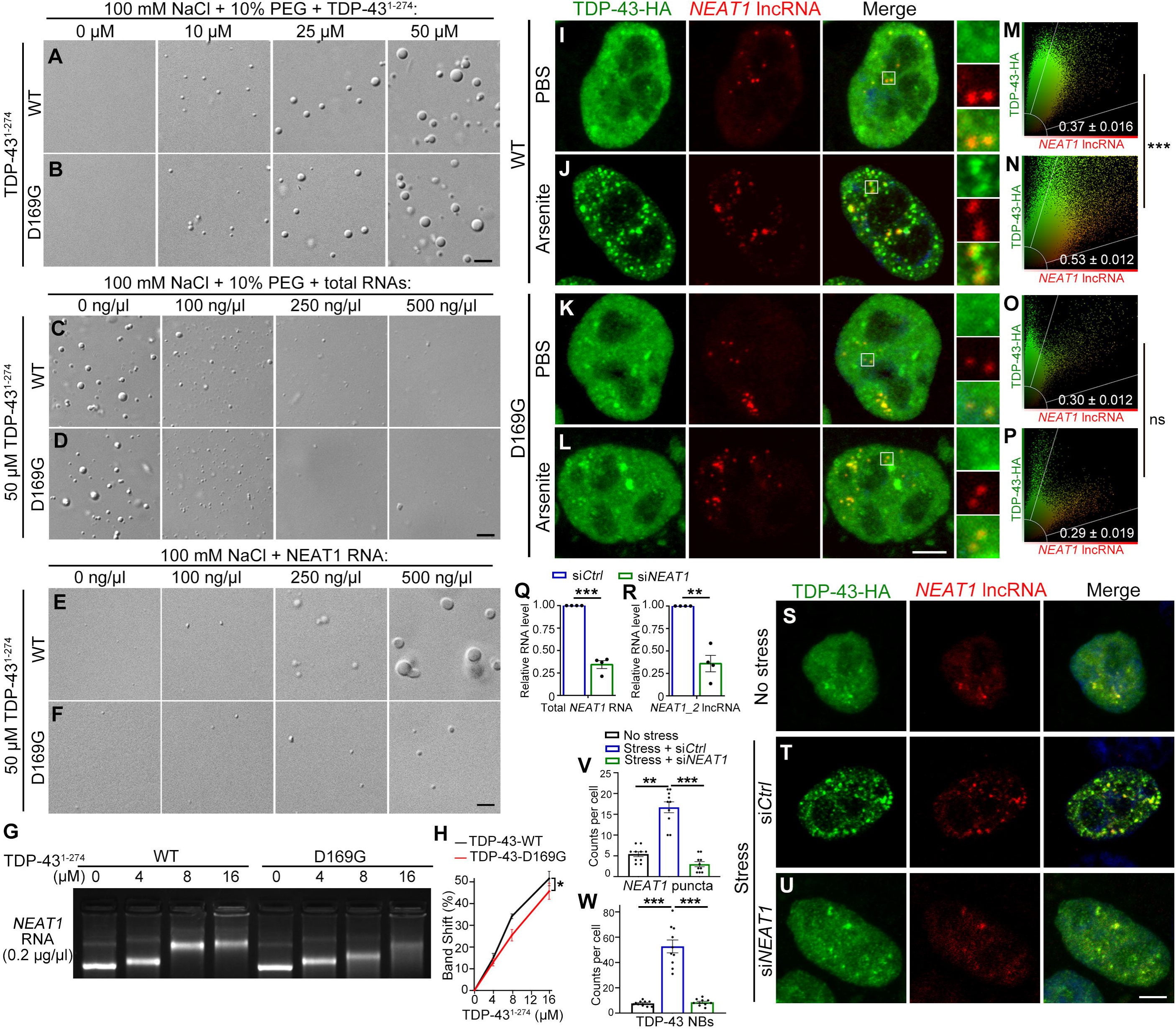
*NEAT1*-mediated nucleation of TDP-43 droplets *in vitro* and in stressed cells is impaired by the D169G mutation. **(A-B)** Both WT and D169G TDP-43 form LDs by LLPS in a dose-dependent manner. **(C-D)** Suppression of the *in vitro* LLPS of WT and D169G TDP-43 by total RNA extracts. **(E-F)** Promotion of TDP-43 LLPS by *NEAT1* RNA is dramatically reduced in the D169G TDP-43. The concentrations of NaCl, TDP-43, PEG and RNA used in the *in vitro* assays are as indicated. **(G-H)** The representative EMSA image (G) and quantification (H) of the gel shifts of *NEAT1* RNA (2 ug) incubated with indicated concentrations of WT or D169G TDP-43^1-274^ protein. All shifts are shown as percentage of the distance the control *NEAT1* RNA band migrates (0 µM protein). **(I-L)** Representative confocal images of WT (I-J) or D169G (K-L) TDP-43 NBs induced by arsenite treatment (250 µM, min) compared to the vehicle control (PBS) in HeLa cells. The cells are co-stained for *NEAT1* lncRNA by FISH using a probe that target the middle sequence of *NEAT1_2*. Higher magnification images of the boxed areas are shown on the side. **(M-P)** For the colocalization analysis of the entire cell, the fluorescence intensity of TDP-43-HA and *NEAT1* lncRNA of each pixel in the cells in (I-L) is measured and plotted. Pearson’s correlation coefficient is used as a measure of colocalization. The mean correlation coefficient value ± SEM of ~20 cells per group is shown at the bottom of each representative plot. **(Q-R)** The knockdown efficiency of the siRNA against *NEAT1* (si*NEAT1*) is determined by qPCR. Both total *NEAT1* RNA (Q) and the *NEAT1_2* isoform (R) are examined. **(S-W)** Representative confocal images (S-U) and quantifications (V-W) of stress-induced *NEAT1* puncta (V) or TDP-43 NBs in cells treated with scramble siRNA (siCtrl) or si*NEAT1*. Mean ± SEM, n = 4 in (G, Q-R), n = ~10 cells each group from pooled results of 3 independent repeats in (V-W); \**p* < 0.05, \*\**p* < 0.01, \*\*\**p* < 0.001; ns, not significant; two-way ANOVA in (H), Student’s t-test in (M, R, V, W). Scale bars: 2 µm in (A-F) and 5 µm in (I-L, S-U).

In striking contrast, the induction of TDP-43 LLPS by *NEAT1* RNA was drastically reduced in D169G (Figures 7E and 7F). We then performed the RNA electrophoretic mobility shift assay (EMSA) to evaluate and compare the RNA binding affinity of WT and D169G TDP-43. Incubation of the TDP-43 proteins with *NEAT1* RNA markedly slowed down the migration speed of the latter, which led to a dosage-dependent band shift (Figures 7G). The impact on band shifts caused by D169G was significantly smaller than that of WT TDP-43 (Figures 7G-7H), indicating that the binding affinity of D169G to *NEAT1* is reduced. However, we did notice that at a higher protein concentration (16 µM), the difference between D169G and WT TDP-43 became smaller. This again points tothat the concentrations of nuclear RNAs and proteins are crucial for the assembly-disassembly of TDP-43 NBs, which may partially explain why the stress-induced NB formation of D169G is delayed but not completely abolished (Figures 6C-6E).

To confirm the above *in vitro* findings in the intracellular milieu of mammalian cells, we conducted fluorescence in situ hybridization (FISH) for *NEAT1* lncRNA together with fluorescent immunocytochemistry for WT and D169G TDP-43 in human HeLa cells. We found that stress significantly increased the colocalization of *NEAT1* FISH-RNA foci with WT TDP-43 NBs (Figures 7I-7J, 7M-7N). In contrast, an increase of the colocalization of *NEAT1* lncRNA foci with TDP-43 NBs by stress was not observed with D169G TDP-43 (Figures 7K-7L, 7O-7P), suggesting that *NEAT1* was unable to promote the assembly of D169G TDP-43 NBs in cells *in vivo*. To further confirm the requirement of lncRNA *NEAT1* in the assembly of TDP-43 NBs, we knocked down *NEAT1* by small interfering RNA (siRNA) (Figures 7Q and 7R). Under this condition, the numbers of stress-induced *NEAT1* puncta and TDP-43 NBs are both dramatically reduced (Figures 7S-7W). Together, our findings propose that the paraspeckle scaffolding lncRNA *NEAT1* mediates the assembly of TDP-43 NBs, which may be an important mechanism to mitigate stress and alleviate cytotoxicity, and a compromise of the TDP-43 NB-mediated stress-mitigating mechanism may lead to excessive cytoplasmic translocation and accumulation of ALS-pathogenic, phosphorylated TDP-43 protein inclusions.

## Discussion

### The stress-induced TDP-43 NB and its cytotoxicity-antagonizing role

This study was initiated by the observation of distinct TDP-43 nuclear granules in stressed cells. We then characterized these subnuclear structures and demonstrated that they meet the three criteria of NBs (Stanek and Fox, 2017) – (1) microscopically distinctive; (2) enriched of TDP-43 and partially colocalized with paraspeckles; and (3) highly dynamic and sensitive to the surrounding nucleoplasm. Various cellular stresses such as arsenic stress, nuclear export blockage, and proteasomal inhibition trigger the formation of TDP-43 NBs. And, TDP-43 NBs are observed in stressed human cells, mouse primary neurons, and *Drosophila brains in vivo*. In addition, a recent paper by Gasset-Rosa et al. (2019) showed that TDP-43 at its endogenous level forms nuclear droplets in multiple cell types. Thus, the assembly of TDP-43 NBs occurs in multiple cell types and in different organisms, suggesting a general role and wide participation of TDP-43 NBs in response to cellular stress.

We show in this study that NB-forming TDP-43 such as the NES^mut^ and the K145/192Q mutation are much less cytotoxic than diffused WT TDP-43. More importantly, abolishing the capability of TDP-43-NES^mut^ to form NBs makes the originally non-toxic NES^mut^ exhibit cytotoxicity. How do TDP-43 NBs acquire cytoprotection? It is thought that SGs help cells survive stress by sequestering mRNAs and temporarily arresting protein synthesis (Liu-Yesucevitz et al., 2010). Given that TDP-43 plays an important role in regulating gene expression and RNA metabolism (Tollervey et al., 2010; Han et al., 2012; Seo et al., 2016) and is presented in both SGs and TDP-43 NBs, it is reasonable to speculate that a similar underpinning mechanism may contribute to the cytoprotective function of TDP-43 NBs. For example, the assembly of TDP-43 NBs may stall DNA transcription and/or arrest RNA processing in stressed cells; when stress is relieved, TDP-43 NBs disassemble and release the RNAs and nuclear proteins required for normal cell functions. In addition, our data indicate that TDP-43 NBs can become irreversible protein aggregates under prolonged stress, which is consistent with the observation of intranuclear inclusions of TDP-43 in some ALS cases (Forman et al., 2007).

### The organization of TDP-43 NBs and the functions of the two RRMs

Super-resolution microscopy imaging reveals a core-shell organization of TDP-43 NBs, regulated by the two RRMs differently. TDP-43 contains two canonical RRMs in tandem. RRM1 has a longer Loop3 region than RRM2 and is thought to have a higher affinity for RNA targets (Buratti and Barall, 2001; Kuo et al., 2014). Although ΔRRM1 and ΔRRM2 show similar binding affinity to total RNAs in the dot-blot assay, they do display divergent sensitivities to RNA suppression in the *in vitro* LLPS assays. For example, suppression of TDP-43 LLPS by total RNAs is greatly reduced in ΔRRM1 but only slightly decreased in ΔRRM2. Another example is the assay using tRNA to mimic nucleoplasmic suppression, in which the induction of TDP-43 LLPS by *NEAT1* RNA is much reduced in ΔRRM1 but increased in ΔRRM2.

In cells, WT TDP-43 is diffused in the nucleus under normal conditions and phase separate into NBs when cells are stressed. ΔRRM1 lacks the RRM1-dependent total RNA suppression, leading to spontaneous partition of ΔRRM1 into phase-separated intranuclear droplets. Meanwhile, TDP-43 likely interacts with nucleation factors such as *NEAT1* RNA via the RRM1, which anchors TDP-43 to the core of NBs. As such, without the “centripetal force”, TDP-43-ΔRRM1 is mainly presented in the shell and forms large, ring-shaped subnuclear structures, and stress does not further promote ΔRRM1 to form normal NBs or additional ring-shaped structures. Thus, the RRM1 appears to execute the seemingly conflicting functions – suppressing TDP-43 phase separation under normal conditions whereas condensing TDP-43 to the core of NBs when the LLPS occurs, which might be attained by interacting with different RNAs. In contrast, since ΔRRM2 retains the main RNA suppression mediated by the RRM1, it does not spontaneously phase separate like ΔRRM1. When cells are stressed and the intracellular LLPS of TDP-43 is initiated, ΔRRM2 that lacks the “centrifugal force” to counteract the RRM1-mediated “centripetal force” is condensed to the core and forms small, insoluble NBs. We speculate that the distinct roles of the two RRMs in regulating the assembly of TDP-43 NBs may result from different RNA-recognition patterns and binding preferences. This idea is supported by the later observations in this study that the RRM1 and the RRM2 mediate the promotion of TDP-43 LLPS by *NEAT1* lncRNA and the suppression by tRNA, respectively. Together, we propose a “push-and-pull” model for the distinct functions of the two RRMs in maintaining the “core-shell” organization of TDP-43 NBs.

### The role of lncRNA *NEAT1* in the assembly of TDP-43 NBs

LncRNAs are thought to provide the scaffold for the assembly of NBs such as paraspeckles (Clemson et al., 2009; Chen and Carmichael, 2009; Yamazaki et al., 2018) and nuclear stress bodies (Jolly et al., 2004; Valgardsdottir et al., 2005). In particular, the paraspeckle lncRNA *NEAT1* are increased in patients and/or animal models of several neurodegenerative diseases including ALS (Nishimoto et al., 2013), Alzheimer’s disease (Puthlyedth et al., 2016), Parkinson’s disease (Mariani et al., 2016), Huntington’s disease (Cheng et al., 2018), and multiple sclerosis (Schirmer et al., 2019).

In this study, we find that *NEAT1* lncRNA is substantially upregulated in stressed neurons and TDP-43 NBs are partially colocalized with *NEAT1* lncRNA foci. In the meanwhile, knockdown of *NEAT1* substantially reduces the formation of stress-induced TDP-43 NBs. Further *in vitro* experiments indicate that *NEAT1* not only promotes TDP-43 LLPS, but also antagonizes the “suppressive” environment modeled with tRNA. As *NEAT1_2* is a lncRNA over 20 kb in length and has complex secondary structures, we speculate that it may provide multiple binding sites for gathering TDP-43 molecules (and potentially other *NEAT1*-binding proteins) together. This would increase the multivalent interactions leading to co-phase separation of TDP-43 and *NEAT1*. In contrast, shorter RNAs such as tRNA lacking long stretches to accommodate multiple TDP-43 molecules may instead segregate each individual TDP-43, therefore suppressing the phase separation. As such, stress-induced upregulation of *NEAT1* may provide nucleation scaffolds to concentrate TDP-43 and other NB proteins, overturning the suppressive environment in the nucleoplasm, leading to the emergence of TDP-43 NBs. It is worth noting that lncRNAs such as *NEAT1* are RNA targets of TDP-43 (Polymenidou et al., 2011; Chung et al., 2018; Modic et al., 2019), implying that complicated cross-regulations between TDP-43 and *NEAT1* as well as between SGs and paraspeckles (An et al. 2019) might be involved in stress and diseases.

### The D169G mutation and the relevance of TDP-43 NBs in ALS pathogenesis

More than disease-causing mutations have been identified in TDP-43, most of which are in the LCD and the adjacent regions (Chen-Plotkin et al., 2010; Lee et al., 2011). D169G is the only known ALS-causing mutation within the RRM1 of TDP-43 (Kabashi et al., 2008) and its pathogenic mechanism has been elusive. Initially, it was presumed to abrogate RNA binding due to its location within the RRM (Kabashi et al., 2008). However, subsequent work indicates that the mutation by D169G causes a small local conformational change in the Turn6 of the β-sheets within the RRM1 of TDP-43 (Austin et al., 2014; Chiang et al., 2016), but does not reduce the overall binding affinity of TDP-43 to RNA or DNA (Kuo et al., 2014). And, although D169G exhibits slightly increased thermal stability and may be more susceptible to proteolytic cleavage (Austin et al., 2014; Chiang et al., 2016), it does not engender aberrant oligomerization but rather increases the resistance of TDP-43 to aggregation (Austin et al., 2014). The only known major cellular alteration by D169G is that it reduces polyubiquitination and the co-aggregation of TDP-43 with Ubiquilin 1 (Kim et al., 2009). However, given the long-standing notion that cytoplasmic inclusions containing ubiquitinated TDP-43 is a pathological hallmark and cause of the disease (Li et al., 2013; Jovičić et al., 2016; Zhao et al., 2018), it has been difficult to understand how decreased ubiquitination and aggregation of TDP-43-D169G causes ALS.

In this study, we discover that D169G has a striking and specific deficit in stress-induced TDP-43 NBs. In line with that, the induction of TDP-43 LLPS by *NEAT1* is dramatically diminished in the *in vitro* assay and the co-localization of TDP-43 NBs with *NEAT1* lncRNA foci is also reduced in D169G. In contrast, the suppression of TDP-43 LLPS by total RNAs is unaffected by D169G and the recruitment of D169G to SGs is not impeded. Instead, the NB-forming defective D169G forms significantly more TDP-43^+^ SGs when cells are subjected to prolonged cellular stress, which gives rise to cytoplasmic TDP-43 foci that are marked with the phosphorylation disease hallmark pS409/410. Together, we propose that the stress-induced upregulation of lncRNA *NEAT1* promotes the assembly of TDP-43 NBs via LLPS, which may be called up to engage in the first line of defense against cellular stresses and disease conditions. Prompt assembly of TDP-43 NBs may lower the need for forming cytoplasmic SGs and prevents excessive cytoplasmic translocation and accumulation of TDP-43. The ALS-causing D169G mutation provides a paradigm for compromise of TDP-43 NBs leading to abnormal cytoplasmic foci containing pTDP-43, which may contribute to ALS pathogenesis. Given that cytoplasmic mislocalization and nuclear depletion of TDP-43 are common in diseased neurons, it is likely that loss of the TDP-43 NB-mediated stress-mitigating function may also underlie other cases of ALS and related diseases.

## Acknowledgments

We thank the BDSC for providing the fly strains, the SIBCB *Drosophila* Center for the fly embryo injection services, L. Chen, L. Pan, J. Zhou, and Z. Zhang for the cloning vectors and plasmids, Z. Zhang for help in the pulse-chase assay, S. Qiu for technical supports, J. Yuan, A. Li, Y. Chen and Z. He for comments and critical reading of the manuscript, and the members of the Fang lab for helpful discussion.

## Author contributions

G.D., Z.M. and Y.F. conceived the research; C.W., Y.D., G.D., C.L., L.G. and Y.F. designed the experiments; C.W., Y.D. G.D., Q.W., K.Z., X.D., B.Q., J.G. and S.Z. performed the experiments; C.W., Y.D., G.D., Q.W., Z.M., J.G. and C.L contributed important new reagents; C.W., Y.D. G.D., and Y.F. analyzed the data and interpreted the results; C.W., Y.D., G.D. and Y.F. prepared the figures; and C.W., Y.D. G.D., C.L. and Y.F. wrote the paper. All authors read and approved the final manuscript. Competing interests: None. Data and materials availability: All data are available in the manuscript or the supplementary materials.

## Supplemental Information

Online Methods

Figures S1-S7

Videos S1-S2

