## Supplemental Information for "Stress induces dynamic, cytotoxicity-antagonizing TDP-43 nuclear bodies via paraspeckle lncRNA *NEAT1*-mediated liquid-liquid phase separation"

### Supplemental Inventory

**1. Online Methods**

**2. Supplemental Figures**

Figure S1, related to Figure 1

Figure S2, related to Figure 2 and Figure 3

Figure S3, related to Figure 3 and Figure 5

Figure S4, related to Figure 3

Figure S5, related to Figure 4

Figure S6, related to Figure 5

Figure S7, related to Figure 6 and Figure 7

**3. Supplemental Videos**

Movie S1, related to Figure 1

Movie S2, related to Figure 1

### Online Methods

#### Plasmids and constructs

The pcDNA3.1-TDP-43-HA and pCMV-myc-TDP-43 plasmids were as previously described (Sun et al., 2018) and used as the templates to generate the following plasmids. The pcDNA3.1-TDP-43-NES$^{mut}$, NLS$^{mut}$, D169G ,Q331K and M337V-HA plasmids were generated by site-directed mutagenesis using the Fast Mutagenesis Kit II (Vazyme). The pcDNA3.1-TDP-43-K145/192Q-HA and pcDNA3.1-TDP-43-NES$^{mut}$-K145/192R-HA plasmids were generated by site-directed mutagenesis using the Fast Mutagenesis Kit MultiS (Vazyme). The pCMV-myc-TDP-43$^{1\text{-}274}$ pCMV-myc-TDP-43-ΔRRM1, and pCMV-myc-TDP-43-ΔRRM2 plasmids were generated by PCR using the pCMV-myc-TDP-43 plasmid as a template. The pcDNA3.1-TDP43-ΔRRM1-HA and pcDNA3.1-TDP43-ΔRRM2-HA plasmids were generated by PCR using the pCMV-myc-TDP-43-ΔRRM1 and pCMV-myc-TDP-43-ΔRRM2 plasmids as a template. To generate the pCAG-GFP-TDP-43 plasmid, the GFP coding sequence was amplified from pcDNA3.1-GFP-AXR3 (a gift from Dr. Z. Zhang) by PCR and then subcloned into a pCAG-hTDP-43 plasmid (Sun et al., 2018) using the ClonExpress MultiS One Step Cloning Kit (Vazyme). The pCAG-GFP-TDP-43-NES$^{mut}$ plasmid was generated by site-directed mutagenesis using the Fast Mutagenesis Kit II (Vazyme). To generate the pCAG-GFP-SFPQ plasmid, The SFPQ coding sequence was amplified from pmEmerald-C1-SFPQ (a gift from Dr. L. Chen) by PCR and then subcloned into the pCAG-GFP-TDP-43 plasmid using the ClonExpress MultiS One Step Cloning Kit (Vazyme).

For generation of transgenic fly lines, WT and various mutant UAS-hTDP-43 constructs were generated by PCR using the above plasmids as a template and sub-cloned into the pBID-UASC-6*Myc vector (Cao et al., 2017) between the KpnI and ApaI sites.

For lentivirus infection of mouse primary neurons, the pLenti-hSyn-TDP-43-HA, pLenti-hSyn-TDP-43-ΔRRM1-HA and pLenti-hSyn-TDP-43-ΔRRM2-HA plasmid was generated by PCR using the pcDNA3.1-TDP-43-HA as a template and sub-cloned into the pLenti-hSyn vector (a gift from Dr. Y. Chen).

For *Escherichia coli* (*E. coli*) expression, the pET28a-6×His-sumo-TDP-43, His-sumo-TDP-43-$^{1-274}$, His-sumo-TDP-43-ΔRRM1, His-sumo-TDP-43-ΔRRM2, His-TDP-43$^{1-274}$, His-TDP-43$^{1-274}$-ΔRRM1, His-TDP-43$^{1-274}$-ΔRRM2, His-TDP-43$^{1-274}$-K145/192Q, His-TDP-43$^{1-274}$-NES$^{mut}$ and His-TDP-43$^{1-274}$-NES$^{mut}$-K145/192R and His-TDP-43$^{1-274}$-D169G constructs were generated by PCR using the above WT or mutant TDP-43 plasmid as a template and were sub-cloned into a pET28a-6×His (a gift from Dr. L. Pan) or pET28a-6×His-sumo (a gift from Dr. J. Zhou) vector. The His-TDP-43$^{1-274}$-K145/192R plasmid was generated by site-directed mutagenesis using the Fast Mutagenesis Kit MultiS (Vazyme) using the pET28a-6×His-TDP-43$^{1-274}$ plasmid a template.

For *in vitro* transcription of NEAT1 RNA, the pcDNA3.1-NEAT1 was generated by PCR using the pEGFP-C1-mNEAT1 plasmid (a gift from Dr. L. Chen) as a template and sub-cloned into the pcDNA3.1 vector.

The primers used for PCR to generate the expression plasmids are summarized below. All constructs were verified by sequencing to ensure the integrity of the cloned open reading frames.

pcDNA3.1-TDP-43-HA:

5'- CGTTTAAACGGGCCCTCTAGAGCCACCATGTCTGAATATATTCGG -3'

5'-CACAGTGGCGGCCGCCTAAGCGTAGTCTGGGACGTCGTATGGGTACATTCCCCAG CCAGAAG -3'

pcDNA3.1-TDP-43-NLS$^{mut}$-HA:

5'- TAACGCAGCAGCAATGGATGAGACAGATGCTTCATCA -3

5'- CCATTGCTGCTGCGTTATCTTTTGGATAGTTGACAACATACA -3

pcDNA3.1-TDP-43-NES$^{mut}$-HA:

5'- GGCAGCGCAGTCTGCATGTGGAGAGGACTTGATCATTAAAGG -3

5'- ATGCAGACTGCGCTGCCTGATCATCTGCAAATGTAACAAAGG -3

pLenti-hSyn-TDP-43-HA:

5'- AGAGCGCAGTCGAGAGGATCCGCCACCATGTCTGAATATATTCGG -3'

5'- GATAAGCTTGATATCGAATTCTCATTAAGCGTAGTCTGGGACGTCG -3'

pLenti-hSyn-TDP-43-ΔRRM1-HA:

5’- AGAGCGCAGTCGAGAGGATCCGCCACCATGTCTGAATATATTCGG -3’

5’- GATAAGCTTGATATCGAATTCTCATTAAGCGTAGTCTGGGACGTCG -3’

pLenti-hSyn-TDP-43-ΔRRM2-HA:

5’- AGAGCGCAGTCGAGAGGATCCGCCACCATGTCTGAATATATTCGG -3’

5’- GATAAGCTTGATATCGAATTCTCATTAAGCGTAGTCTGGGACGTCG -3’

pBID-UASC-TDP-43-Myc:

5’- CGGCCGCGGCTCGAGGGTACCATGTCTGAATATATTCGGGTAACCG -3’

5’- GAGTTTTTGTTCGAAGGGCCCTCTAGACTCGAGCATTCCCCAGCCAGAAGACTT-3’

pBID-UASC-TDP-43-NLS$^{mut}$-Myc:

5’- CGGCCGCGGCTCGAGGGTACCATGTCTGAATATATTCGGGTAACCG -3’

5’- GAGTTTTTGTTCGAAGGGCCCTCTAGACTCGAGCATTCCCCAGCCAGAAGACTT-3’

pBID-UASC-TDP-43-NES$^{mut}$-Myc:

5’- CGGCCGCGGCTCGAGGGTACCATGTCTGAATATATTCGGGTAACCG -3’

5’- GAGTTTTTGTTCGAAGGGCCCTCTAGACTCGAGCATTCCCCAGCCAGAAGACTT-3’

pcDNA3.1-TDP-43-K145/192Q-HA:

5’- TCATTCACAGGGGTTTGGCTTTGTTCGTTTT -3

5’- CAAACCCCTGTGAATGACCAGTCTTAAGATCTTTCTTG -3

5’- GCAGACAAGTGTTTGTGGGGCGCTGTAC -3

5’- CACAAACACTTGTCTGCTTCTCAAAGGCTCATCTT -3

pcDNA3.1-TDP-43-NES$^{mut}$-K145/192R-HA:

5’- TCATTCAAGGGGGTTTGGCTTTGTTCGTTTT -3

5’- CAAACCCCCTTGAATGACCAGTCTTAAGATCTTTCTTG -3

5’- GCAGAAGAGTGTTTGTGGGGCGCTGTAC -3

5’- CACAAACACTCTTCTGCTTCTCAAAGGCTCATCTT -3

pcDNA3.1-TDP-43-D169G-HA:

5’- ATGATAGGTGGACGATGGTGTGACTGCAAACT -3’

5’- CATCGTCCACCTATCATATGTCGCTGTGACATTACTTTC -3’

pcDNA3.1-TDP-43-Q331K-HA:

5’- CAGCACTAAAGAGCAGTTGGGGTATGATGGGC -3’

5’- ACTGCTCTTTAGTGCTGCCTGGGCGGCAGCCA -3’

pcDNA3.1-TDP-43-M337V-HA:

5’- GGTATGGTGGGCATGTTAGCCAGCCAGCAGAA -3’

5’- AACATGCCCACCATACCCCAACTGCTCTGTAGTGC -3’

pCMV-Myc-TDP-43:

5’- ATGGCCATGGAGGCCCGAATTCATGTCTGAATATATT-3’

5’- CCGCGGCCGCGGTACCTCGAGCTACATTCCCCAGCCAGAAGAC -3’

pCMV-Myc-TDP-43$^{1–274}$:

5’- ATGGCCATGGAGGCCCGAATTCATGTCTGAATATATT -3’

5’- CCGCGGCCGCGGTACCTCGAGCTATCCACTTCTTTCTAACTGTCTATTGC -3’

pCMV-Myc-TDP-43-ΔRRM1:

5’- ATGGCCATGGAGGCCCGAATTCATGTCTGAATATATT -3’

5‘- CCGCGGCCGCGGTACCTCGAGCTACATTCCCCAGCCAGAAGAC -3’

pCMV-Myc-TDP-43-ΔRRM2:

5’- ATGGCCATGGAGGCCCGAATTCATGTCTGAATATATT -3’

5‘- CCGCGGCCGCGGTACCTCGAGCTACATTCCCCAGCCAGAAGAC -3’

pcDNA3.1-TDP43-ΔRRM1-HA:

5’- CGTTTAAACGGGCCCTCTAGAGCCACCATGTCTGAATATATTCGG -3’

5’-ATATCCAGCACAGTGGCGGCCGCTTAAGCGTAGTCTGGGACGTCGTATGGGTACAT TCCCCAGCCAGAAGACTTA -3’

pcDNA3.1-TDP43-ΔRRM2-HA:

5’- CGTTTAAACGGGCCCTCTAGAGCCACCATGTCTGAATATATTCGG -3’

5’-ATATCCAGCACAGTGGCGGCCGCTTAAGCGTAGTCTGGGACGTCGTATGGGTACAT TCCCCAGCCAGAAGACTTA -3’ pCAG-GFP-TDP-43:

5’- CATCATTTTGGCAAAGAATTCCACCATGGTGAGCAAGGGCGAGG -3’

5’- CAGACATGCTTCCGCCCTTGTACAGCTCGTCCATGCC -3’

5’- CGAGCTGTACAAGGGCGGAAGCATGTCTGAATATATTCGGGTAACCG -3’

5’- GCTCCCCGGGGGTACCTCGAGCTATTACATTCCCCAGCCAGAAGACTT -3’

pCAG-GFP-TDP-43-NES$^{mut}$:

5’- GGCAGCGCAGTCTGCATGTGGAGAGGACTTGATCATTAAAGG -3

5’- ATGCAGACTGCGCTGCCTGATCATCTGCAAATGTAACAAAGG -3

pCAG-GFP-SFPQ:

5’- CATCATTTTGGCAAAGAATTCCACCATGGTGAGCAAGGGC -3

5’- AGACATGCTTCCGCCCTTGTACAGC -3

5’- ACAAGGGCGGAAGCATGTCTCGGGATCGGTTCCG -3

5’- GCTCCCCGGGGGTACCTCGAGCTAAAATCGGGGTTTTTTGTTTG -3

pET28a-6×His-TDP-43$^{1–274}$:

5’- CAGCAAATGGGTCGCGGATCCATGTCTGAATATATTCGGGTAACCG -3’

5’- GTGGTGGTGGTGGTGCTCGAGTTAACTTCTTTCTAACTGTCTATTGCTATTG -3’

pET28a-6×His-TDP-43$^{1–274}$-ΔRRM1:

5’- CAGCAAATGGGTCGCGGATCCATGTCTGAATATATTCGGGTAACCG -3’

5’- GTGGTGGTGGTGGTGCTCGAGTTAACTTCTTTCTAACTGTCTATTGCTATTG -3’

pET28a-6×His-TDP-43$^{1–274}$-ΔRRM2:

5’- CAGCAAATGGGTCGCGGATCCATGTCTGAATATATTCGGGTAACCG -3’

5’- GTGGTGGTGGTGGTGCTCGAGTTAACTTCTTTCTAACTGTCTATTGCTATTG -3’

pET28a-6×His-TDP-43$^{1–274}$-K145/192Q:

5’- CAGCAAATGGGTCGCGGATCCATGTCTGAATATATTCGGGTAACCG -3’

5’- GTGGTGGTGGTGGTGCTCGAGTTAACTTCTTTCTAACTGTCTATTGCTATTG -3’

pET28a-6×His-TDP-43$^{1–274}$-K145/192R:

5’- TCATTCAAGGGGGTTTGGCTTTGTTCGTTTT -3

5’- CAAACCCCCTTGAATGACCAGTCTTAAGATCTTTCTTG -3

5’- GCAGAAGAGTGTTTGTGGGGCGCTGTAC -3

5’- CACAAACACTCTTCTGCTTCTCAAAGGCTCATCTT -3

pET28a-6×His-TDP-43$^{1–274}$-NES$^{mut}$:

5’- CAGCAAATGGGTCGCGGATCCATGTCTGAATATATTCGGGTAACCG -3’

5’- GTGGTGGTGGTGGTGCTCGAGTTAACTTCTTTCTAACTGTCTATTGCTATTG -3’

pET28a-6×His-TDP-43$^{1-274}$-NES$^{mut}$-K145/192R:

5’- CAGCAAATGGGTCGCGGATCCATGTCTGAATATATTCGGGTAACCG -3’

5’- GTGGTGGTGGTGGTGCTCGAGTTAACTTCTTTCTAACTGTCTATTGCTATTG -3’

pET28a-6×His-TDP-43$^{1–274}$-D169G:

5’- CAGCAAATGGGTCGCGGATCCATGTCTGAATATATTCGGGTAACCG -3’

5’- GTGGTGGTGGTGGTGCTCGAGTTAACTTCTTTCTAACTGTCTATTGCTATTG -3’

pET28a-6×His-sumo-TDP-43:

5’- AGAGAACAGATTGGTGGATCCATGTCTGAATATATTCGGGTAACCG -3’

5’-TTGTCGACGGAGCTCGAATTCTTACATTCCCCAGCCAGAAGAC -3’

pET28a-6×His-sumo-TDP-43-$^{1–274}$:

5’- AGAGAACAGATTGGTGGATCCATGTCTGAATATATTCGGGTAACCG -3’

5’- TTGTCGACGGAGCTCGAATTCTTATCCACTTCTTTCTAACTGTCTATTGC -3’

pET28a-6×His-sumo-ΔRRM1:

5’- AGAGAACAGATTGGTGGATCCATGTCTGAATATATTCGGGTAACCG -3’

5’-TTGTCGACGGAGCTCGAATTCTTACATTCCCCAGCCAGAAGAC -3’

pET28a-6×His-sumo-ΔRRM2:

5’- AGAGAACAGATTGGTGGATCCATGTCTGAATATATTCGGGTAACCG -3’

5’-TTGTCGACGGAGCTCGAATTCTTACATTCCCCAGCCAGAAGAC -3’

pcDNA3.1-NEAT1:

5’- CGTTTAAACGGGCCCTCTAGAGAGTTAGTGACAAGGAGGGCTCG -3’

5’- ATATCCAGCACAGTGGCGGCCGCTTCAATCTCAAACCTTTATTTTGCTG-3’

**Cell cultures and transfection**

293T and HeLa cells were cultured in Dulbecco's Modified Eagle Medium (Sigma, D0819) supplemented with 10% (v/v) fetal bovine serum (FBS, BioWest) and 1% penicillin/streptomycin at 37˚C in 5% $CO_2$. Transient transfection was performed using Lipofectamine™ 3000 (Invitrogen) in Opti-MEM (Invitrogen). Cells were transfected for at least 24h before the subsequent drug treatments or examinations. For the knockdown experiment, the siRNA (Genepharma) was transfect into the HeLa cells using the Lipofectamine™ RNAiMax Transfection Reagent (Invitrogen) according to the manufacturer's instruction. The siRNA was incubated for ~48 h before cells were harvested. The siRNA oligos used in this study are listed below:

si*Ctrl*: 5'- UUCUCCGAACGUGUCACGUTT -3'

si*NEAT1*: 5'- UUACAAAAUAUGUUGCCAUTT -3'

**Pharmacological experiments:**

Arsenite treatment: HeLa or 293T cells were grown on coverslips in a 24-well plate and transfected with the indicated plasmids for 24 h. Cells were then treated with 250 μM of $NaAsO_2$ or PBS for 30 min, prior to fixation with 4% paraformaldehyde. For the recovery experiments, the culture medium containing $NaAsO_2$ was removed and the cells were incubated in fresh medium for indicated time prior to fixation.

LMB treatment: For the nuclear export inhibition assays, LMB was added into the culture medium after 6h transfection at indicated final concentrations.

CHX treatment: For the pulse-chase assays, CHX was added into the medium at each time point at a final concentration of 25 ng/ml.

MG132 treatment: For the proteasomal inhibition assays, MG132 was added into the medium at each time at a final concentration of 25 μM.

CQ treatment: For autophagy inhibition assays, CQ was added into the medium at each time at a final concentration of 25 mM.

The cells incubated in the culture medium with the above drugs for indicated time before fixation for immunocytochemistry or western blotting analysis.

**Immunocytochemistry and immunohistochemistry assays**

HeLa or 293T cells grown on coverslips pre-coated with PLL (Sigma) in a 24-well plate were transfected and treated as described above. The cells were then fixed in 4% paraformaldehyde in PBS for 15 min at room temperature (RT), permeabilized in 0.5% Triton X-100 (Sigma) in PBS for 15 min and blocked with 3% goat serum in PBST (0.1% Triton X-100 in PBS) for 1h at RT. The above primary and secondary antibodies were then incubated in the blocking buffer at 4°C overnight or at RT for 1-2 h. After 3 washes with PBST, cells were mounted on glass slides using the VECTASHIELD Antifade Mounting Medium with DAPI (Vector Laboratories).

**Confocal and super-resolution imaging**

Fluorescent confocal images were taken with Leica TCS SP8 confocal microscopy system using a 63X or 100X oil objective (NA=1.4). Super-resolution images were captured using the Leica SP8 LIGHTNING confocal microscope, which allowed simultaneous multicolor imaging in super-resolution down to 120 nm. Confocal or super-resolution images were then processed in LAS X (Leica) and assembled into figures using Adobe Photoshop CS6.

**Live cell imaging**

HeLa cells transfected with pCAG-GFP-TDP-43 were grown on Nunc™ Lab-Tek™ Chambered Coverglass (Thermo) for 24 h before the cells were treated with 250 μM of $NaAsO_2$. For live cell imaging, the chambered coverglasses were placed in the Incubation System for Microscopes (Tokai Hit) and maintained at 37˚C in 5% $CO_2$ for the duration of the experiment. Time-lapse images were captured using the Leica TCS SP8 confocal microscopy system every 10 min for about 1 h.

### Fluorescence recovery after photobleaching (FRAP) assay

The PFRP assay was performed using the FRAP module of the Leica SP8 confocal microscopy system. In brief, each GFP-TDP-43 NB was bleached using a 488 nm laser at 100% laser power for approximately 5 s. After photobleaching, time-lapse images were captured every 10 s for the about 5 min. For each indicated time point (t), the fluorescence intensity within the bleached NB was normalized to the fluorescence intensity of a nearby, unbleached NB (to control for photobleaching during prolonged live imaging). The normalized fluorescence intensity of pre-bleaching was set to 100% and the normalized fluorescence intensity at each time point ($I_t$) was used to calculate the fluorescence recovery according to the following formula: FR(t) = $I_t/I_{pre\text{-}bleaching}$. Image J was used for quantification and GraphPad Prism to plot and analyze the FRAP experiments.

### Antibodies

The following antibodies were used for Western blotting, immunoprecipitation and immunofluorescence assays: mouse anti-FLAG (Sigma, F3165), mouse anti-HA (Proteintech, 66006-1), mouse anti-pS409/410-TDP-43 (Proteintech, 66318-1-Ig), mouse anti-p54/nrb NONO (Santa Cruz, sc-166702), mouse anti-G3BP (BD Biosciences, 611127), rabbit anti-HA (CST, C29F4), rabbit anti-c-Myc (Sigma, c3956), rabbit anti-TDP-43 (Proteintech, 10782-2-AP), anti-β-Tubulin III (Sigma, T2200), rabbit anti-TIAR (Cell Signaling Technology, 8509S), rabbit anti-SC35 (Abcam, ab204916), rabbit anti-SFPQ (Abcam, ab177149), rabbit anti-TAF9 (Abcam, ab169784), rabbit anti-PML (Abcam, ab179466), and chicken anti-MAP2 (Abcam, ab5392). HRP conjugated secondary antibodies: goat anti-mouse (Sigma, A4416) and goat anti-rabbit (Sigma, A9169). Fluorescent secondary antibodies: goat anti-mouse-Alexa Fluor 488 (Life Technologies, A11029), goat anti-rabbit-Alexa Fluor 568 (Life Technologies, A11036) and goat anti-Chicken-Alexa Fluor 568 (Life Technologies, A11041).

**Protein extraction and Western blotting**

Total protein was extracted from cells in a 2% SDS extraction buffer (50 mM Tris pH 6.8, 2% SDS, 1% mercaptoethanol, 12.5% glycerol and 0.04% bromophenol blue) containing the protease inhibitor cocktail (Roche, 04693132001). For separation of soluble and insoluble proteins, cells or fly heads were lysed on ice in a RIPA buffer (50 mM Tris pH 8.0, 150 mM NaCl, 1% NP-40, 5 mM EDTA, 0.5% sodium deoxycholate, 0.1% SDS) supplemented with protease and phosphatase inhibitors (Roche). After sonication, the homogenates were centrifuged at 16,000 g for 15min at 4°C. The supernatant was used as the soluble fraction and the pellets containing the insoluble fraction were dissolved in a urea buffer (9 M urea, 50 mM Tris buffer, pH 8.0) after wash.

All protein samples were then boiled at 100°C for 5 min and separated using a 10% Bis-Tris SDS-PAGE (Invitrogen). Detection was performed using the High-sig ECL Western Blotting Substrate (Tanon). Images were captured using an Amersham Imager 600 (GE Healthcare) and densitometry was measured using ImageQuant TL Software (GE Healthcare). The contrast and brightness were optimized equally using Adobe Photoshop CS6. All experiments were normalized to tubulin or GAPDH as indicated in the figures.

**Cell viability assay**

Transfected 293T cells were seeded in 96-well plates (Corning) at the density of $9\times10^3$ cells/well and cultured in 100 μL of culture medium. Otherwise, cell viability was examined 48-72 h after transfection using the Cell Counting Kit-8 (CCK-8) (Dojindo), according to the manufacturer's instructions. Briefly, 10 μL of the CCK-8 solution were added to each well and incubated at 37°C for 2.5 h. Finally, the absorbance at 450 nm was measured with a Synergy2 microplate reader (BioTek Instruments).

**ATP level measurement**

Transfected 293T cells were seeded in 96-well plates (Corning) at the density of $2.5\times10^4$

cells/well. ATP levels were examined 36 h after transfection using the CellTiter-Glo® Luminescent Assay (Promega) according to the manufacturer's instructions. Briefly, the CellTiter-Glo® reagent was added into the microplate wells, mixed and incubated for 15 min at 37°C on the shaker. The luminescence was then measured with a Synergy2 microplate reader (BioTek Instruments).

#### *Drosophila* strains

The transgenic fly strains of WT and various mutant UAS-*hTDP-43* were generated by ΦC31 integrase-mediated, site-specific integration, which allowed uniform transgene expression across different lines. The attP2 landing site stock used for the fly embryo injection and transformation in this study was y[1] M{vas-int.Dm}ZH-2A w[*]; P{CaryP}attP40 (25C6). The pBID-UASC-Luciferase (UAS-Luc) transgenic fly strain was generated using the same approach at the same landing site (Cao et al., 2017) and was therefore used as a control in this study. The following strains were obtained from the Bloomington *Drosophila* Stock Center (BDSC): *D42*-Gal4 (#8816), *elav*-Gal4 (#8760) and UAS-*LacZ* (#8529). Flies tested in this study were raised on standard cornmeal media and maintained at 25 °C and 60% relative humidity.

#### Eye degeneration analysis and climbing assay in flies

To assess the eye degeneration, z-stack images of adult fly eyes were captured using an Olympus SZX16 stereomicroscope at indicated ages. Degeneration was evident and assessed by rough surface, swelling and loss of pigment cells of the compound eyes. Each fly eye was single-blindly scored in a scale of 0 to 4, with 0 for no degeneration and 4 for the full degeneration.

For the climbing assay, ~20 flies per vial, 5~10 vials per group were tested. All flies were transferred into an empty polystyrene vial and allowed 15 min for flies to recover. The flies were then gently tapped down to the bottom of the vial and the number of flies that climbed over 3 cm within 10 seconds was recorded. The test was repeated three times for each vial and the average was used in the quantifications.

**Mouse care and surgical procedures**

All mouse procedures were performed in compliance with the institutional guidelines on the scientific use of living animals at Interdisciplinary Research Center on Biology and Chemistry, the Chinese Academy of Sciences (CAS). “Principles of laboratory animal care" (NIH publication No. 86-23, revised 1985) were followed. Animal distress and conditions requiring euthanasia were addressed and the number of animals used was minimized.

**Lentivirus production and primary neuron culture**

To generate lentivirus for infecting primary neurons, 293T cells were co-transfected with pLenti-hSyn-TDP-43-HA, psPAX2 and pMD2.G with a ratio of 4:2:1 in Opti-MEM medium using Lipofectamine™ 2000. Culture supernatant was collected at 48 h after transfection and passed through a 0.45-μm filter. Viral particles were concentrated from culture supernatants by Lenti-X™ Concentrator (Clontech). Viral pellets used for neuronal infection were resuspended in Neurobasal medium (Invitrogen).

Primary cortical neurons were isolated from C57BL/6 mouse cortex at embryonic day 17 (E17) and cultured in serum-free Neurobasal medium (Invitrogen) supplemented with 2% B27, GlutaMax, and penicillin-streptomycin (Invitrogen). At 7 days in vitro (DIV), neurons were infected with pLenti-hSyn-TDP-43-HA for 5 days before extraction for RNA or immunofluorescence.

**RNA extraction and real-time quantitative PCR**

For quantitative PCR (qPCR), total RNA was isolated from mouse primary neuron using TRIzol (Invitrogen) according to the manufacturer’s instruction. After DNase (Promega) treatment, the reverse transcription reactions were performed using All-in-One cDNA Synthesis SuperMix kit (Bimake). The cDNA was then used for real-time qPCR using the SYBR Green qPCR Master Mix

(Bimake) with the QuantStudio™ 6 Flex Real-Time PCR system (Life Technologies). The mRNA levels of GAPDH were used as an internal control to normalize the mRNA levels of *NEAT1*. The qPCR primers used in this study are listed below:

Total *mNEAT1*:

5'- ACTCTTGCCCCTCACTCTGA -3'

5'- CAGGGTGTCCTCCACCTTTA -3';

*mNEAT1_2*:

5'- CCCACACCTCAGTGGTTTCT -3'

5'- ACAGAACCAAGGCACAATCC -3';

*mGAPDH*:

5'- CACCATCTTCCAGGAGCGAG -3'

5'- CCTTCTCCATGGTGGTGAAGAC -3';

**Purification of TDP-43 proteins**

WT or mutant TDP-43 protein was expressed in BL21 (DE3) *E. coli* (TransGenBiotech, CD601-03) at 19°C for 16 h after induction by adding 100 μM of IPTG as previously described (*46*). In brief, cells were harvested by centrifugation at 4000 rpm for 20 min at 4 °C and lysed in 50 mL of lysis buffer (50 mM Tris-HCl, 500 mM NaCl, pH 8.0, 10 mM imidazole, 4 mM β-mercaptoethanol, 1 mM PMSF, and 0.1 mg/mL RNase A). After cell lysates were filtered with a 0.22 μm filter, the protein was purified using Ni columns (GE Healthcare, USA) and then eluted in an elution buffer (50 mM Tris-HCl, 500 mM NaCl, pH 8.0, 250 mM imidazole and 4 mM β-mercaptoethanol). The proteins were further purified using the Superdex 200 16/600 columns (GE Healthcare) in a buffer containing 50 mM Tris-HCl pH 7.5, 300 mM NaCl and 2 mM DTT, and freshly frozen in liquid nitrogen and stored at -80 °C. RNase A was routinely added in cell lysates and administrated again before chromatography during the protein purification procedure. All purified proteins were confirmed by Coomassie brilliant blue staining and Western blotting before use.

**Dot-blot binding assay for *in vitro* analysis of RNA-protein interaction**

Purified WT or mutant TDP-43 protein was diluted in a blotting buffer (50 mM Tris-HCl, pH 7.5, 300 mM NaCl, and 5% glycerol) and blotted onto a 0.45 μm nitrocellulose membrane. The membranes were left to dry at room temperature for 30 min and then stained with Ponceau S for 10s. Images were captured using an Amersham Imager 600 (GE Healthcare). After the images were captured using an Amersham Imager 600 (GE Healthcare), the membranes were washed with PBST (0.05% TWEEN 20 in PBS) for 30 min and then incubated in the PBST containing 25 ng/μl total RNAs or NEAT1 RNA for 1h with gentle rocking and rotation at room temperature. The membranes were then washed in PBST and incubated in the SYBR™ Gold Nucleic Acid Gel Stain (Invitrogen) for 10 min. RNAs bound to the membranes were imaged and examined using the Gel Image System (Tanon).

***In vitro* phase separation and RNA buffering assay**

For the *in vitro* LLPS experiments, purified WT or mutant TDP-43 protein was mixed with NaCl at indicated concentrations in a phase separation buffer (50 mM Tris-HCl, pH 7.5 and 5-10% *(*w/v) PEG 8000 (Sigma)) and incubated for 1 min at room temperature. For the RNA buffering assay, the TDP-43 proteins were incubated with total RNAs, tRNA or NEAT1 RNA in the above phase separation buffer with NaCl at indicated concentrations as shown in the figures. Finally, 5 μL of each sample was pipetted onto a coverslip and imaged using a Leica microscope with differential interference contrast (DIC). The total RNAs were extracted from HeLa cells using TRIzol (Invitrogen) according to the manufacturer's instructions. Yeast tRNA was purchased from Invitrogen. The NEAT1 RNA was *in vitro* transcribed and purified using HiScribe™ T7 Quick High Yield RNA Synthesis Kit (NEB). Total RNAs and NEAT1 RNA were used within one day of production.

**The electrophoretic mobility-shift assay (EMSA)**

To examine the RNA binding affinity of WT and D169G TDP-43, the purified TDP-43$^{1-274}$ protein was incubated with *NETA1* RNA in a buffer containing 50 mM Tris-HCl, 500 mM NaCl, pH 8.0 at 37°C for 10 min, and then loaded in 0.65% UltraPure™ Agarose (Thermo) and examined by electrophoresis. The RNA was visualized using the SYBRTM Green II RNA Gel Stain (Invitrogen) in 0.5 X TBE buffer and imaged with UV light using the Gel Image System (Tanon).

**Fluorescence in situ hybridization (FISH)**

The fluorescently-labeled DNA probes used in the FISH assay to detect *NEAT1* lncRNA were synthesized with the Nick Translation DNA Labeling System (Enzo) and Gold 550 dUTP (Enzo). The *NEAT1_2* cDNA templates used in the above reactions were generated from human HeLa cells or mouse NSC-34 cells by reverse transcription reactions (TaKaRa) using the following primers:

Human *NEAT1_2* mid1:

5’-GCCACATTCTTTGCCTTCAT-3’;

5’-TCATTTACCCGCATTTCACA-3’

The FISH assay was conducted as previously described (Mito et al., 2016). Briefly, cells grown on coverslips were fixed in 4% paraformaldehyde (PBS) for 30 min at room temperature (RT), washed twice with PBS, and permeabilized in 0.5% Triton X-100 (Sigma) and 2 mM rnase-inhibitor-ribonucleoside-vanadyl complexes (RVC) (Sangon Biotech) in PBS for 30 min at RT. The hybridization buffer containing the fluorescent labeled DNA probes (10 ng/μl) in 50% Formamide (Sangon Biotech), 2x saline sodium citrate (SSC) (Invitrogen) and 500 μg/ml Salmon Sperm DNA Solution (Invitrogen) was heated at 100 °C for 10 min and placed on ice before incubated with the samples at 37 °C overnight. For the simultaneous detection of TDP-43, immunocytochemistry was performed afterwards as mentioned above.

**Statistical Analysis**

Statistical significance in this study is determined by one-way analysis of variance (ANOVA) with Tukey's HSD post-hoc test, two-way ANOVA with Bonferroni's post-hoc test, or unpaired, two-tailed Student's *t*-test with unequal variance at $*p < 0.05$, $**p < 0.01$, and $***p < 0.001$ as indicated in the legends of each figure. Error bars represent the standard error of the mean (SEM).

### Supplementary Figures

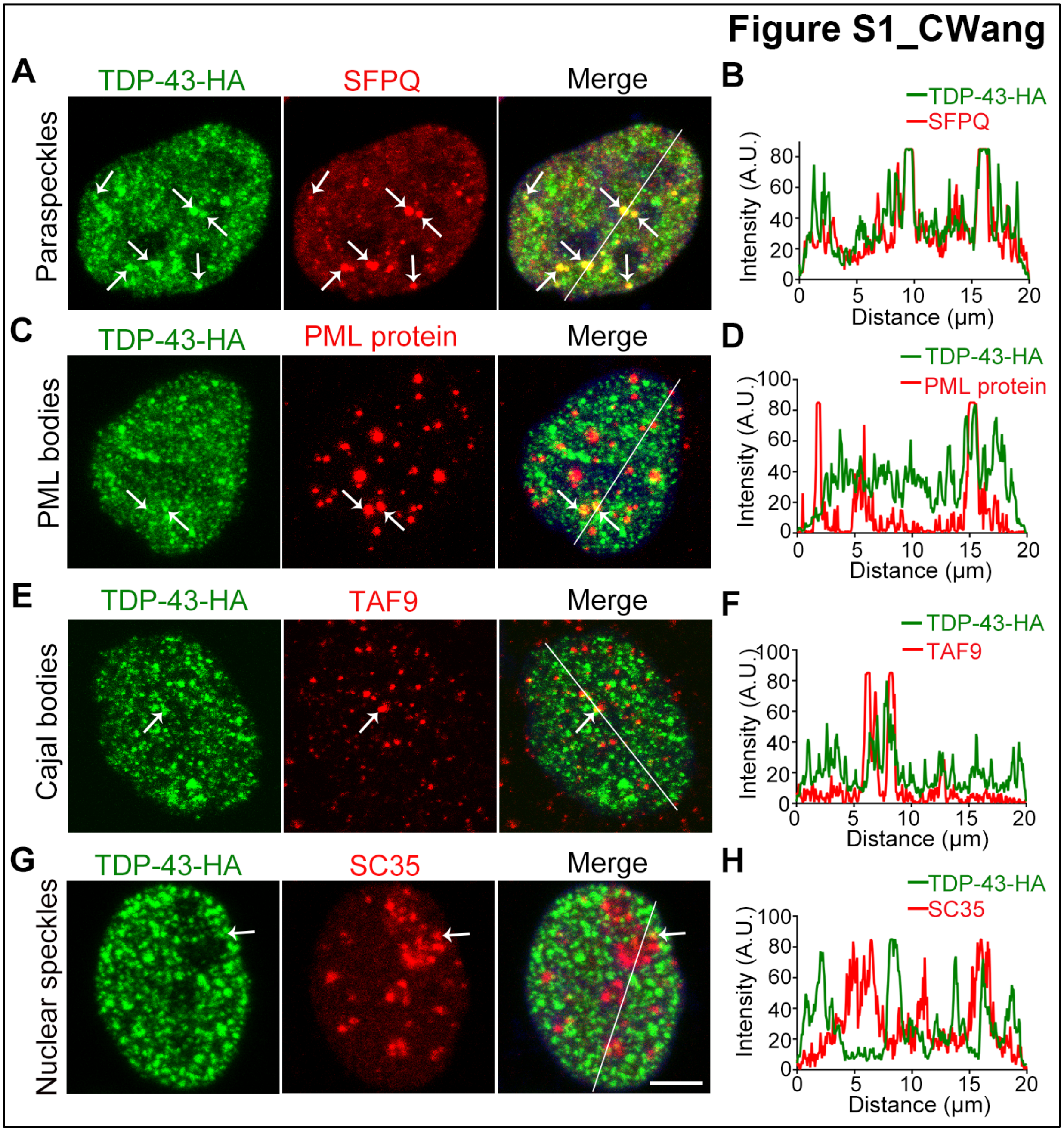


**Figure S1. Co-immunostaining of TDP-43 with several known NB markers, related to Figure 1**

Representative confocal images of the nucleus of 293T cells transfected with TDP-43-HA and treated with 250 μM $NaAsO_2$ for 30 min. All cells are examined by immunocytochemistry with anti-HA for TDP-43-HA (green) and the indicated antibodies (red): (**A-B**) SFPQ (splicing factor proline-glutamine rich), for paraspeckles (Prasanth et al., 2005). (**C-D**) PML (promyelocytic leukaemia) protein, for PML bodies (Koken et al., 1994). (**E-F**) TAF9 (TATA-box binding protein associated factor 9) for Cajal bodies (Santama et al., 2005). (**G-H**) SC35 (serine/arginine-rich splicing factor-like protein), for nuclear speckles (Fu and Maniatis, 1990). Merged images with DAPI staining (for DNA) are shown. Arrows indicate the examples of co-localization of TDP-43 with the indicated NB markers. The co-localization of TDP-43 with each type of NBs is further evaluated by the line scanning analysis shown in (B), (D), (F) and (H). A.U., arbitrary unit. Scale bar: 5 μm. The results indicate that TDP-43 NBs are well colocalized with paraspeckles, and in a few rare cases partial colocalization with PML bodies, Cajal bodies or nuclear speckles are also spotted.

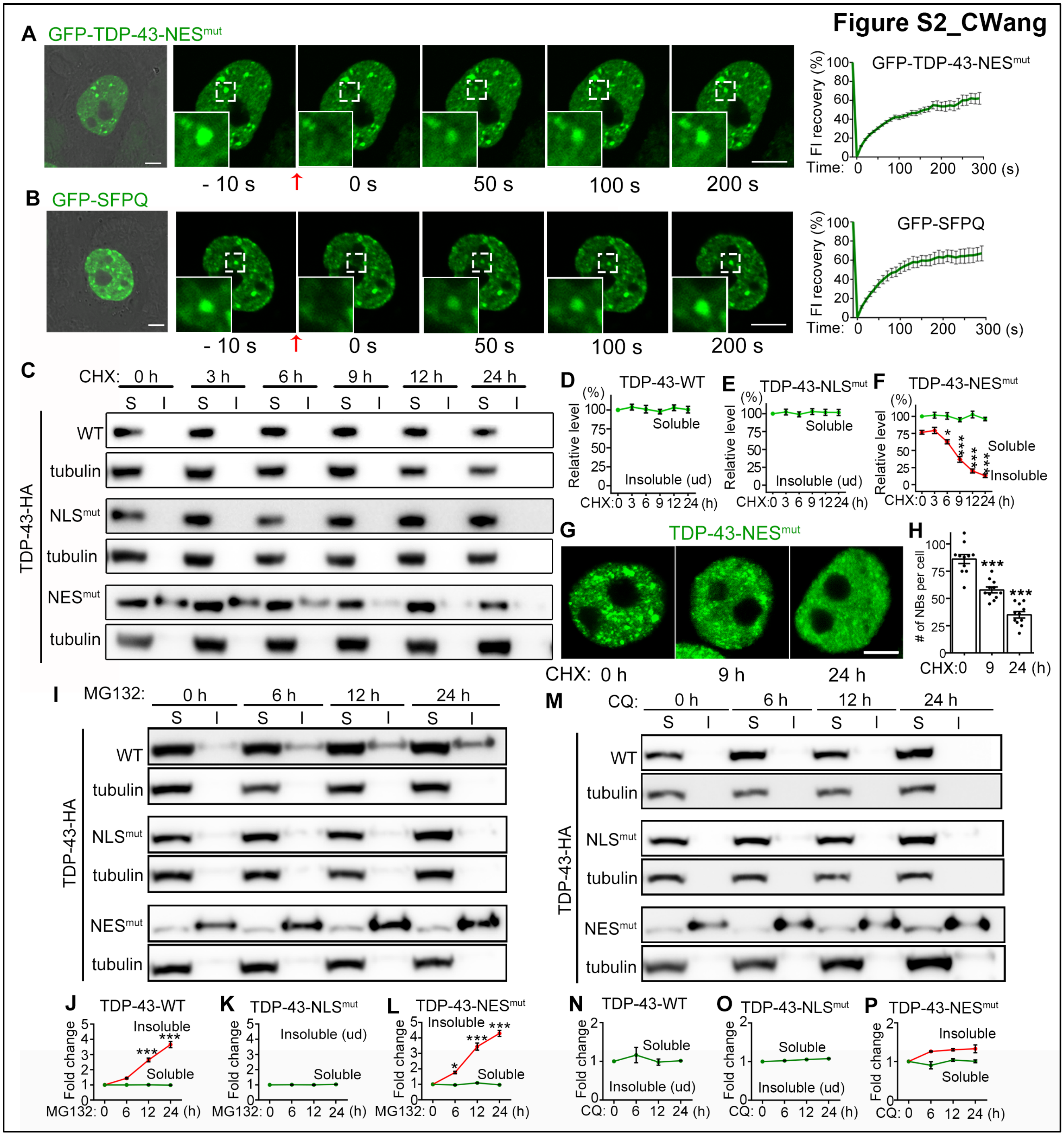


**Figure S2. TDP-43-NES$^{mut}$ forms dynamic and reversible NBs, and the turnover of insoluble TDP-43 is sensitive to proteasomal but not autophagic inhibition, related to Figure 2 and 3**

(**A-B**) Representative images and the recovery curves of the FRAP analysis of the spontaneous NBs of GFP-TDP-43-NES$^{mut}$ (A) or the NBs of GFP-SFPQ induced by arsenite (250 μM, 30 min) (F) in *live* cells. (**C-F**) Representative Western blot images (C) and quantifications (D-F) of the pulse-chase experiments of WT, $NLS^{mut}$ and $NES^{mut}$ TDP-43 protein in the soluble (S, supernatants in RIPA) and the insoluble (I, pellets resuspended in 9 M of urea) fractions at indicated time after Cycloheximide (CHX) treatment. All proteins are normalized to tubulin in the soluble fraction and the relative level at 0 h is set to 100%. (**G**) Representative images showing TDP-43-$NES^{mut}$ NBs in 293T cells at indicated time after CHX treatment. (**H**) Quantification of the average number of TDP-43 NBs per cell in (G). (**I-L**) Representative images (I) and quantifications (J-L) of Western blotting analyses of WT, $NLS^{mut}$ and $NES^{mut}$ TDP-43 in the soluble (S) and insoluble (I) fractions at indicated time after treating the cells with MG132 to inhibit the proteasome-mediated protein degradation. (**M-P**) Representative images (M) and quantifications (N-P) Western blot analyses of WT, $NLS^{mut}$ and $NES^{mut}$ TDP-43 in the soluble (S) and insoluble (I) fractions at indicated time after treating the cells with chloroquine (CQ) to block the autophagic flux. S, supernatants in RIPA; I, precipitates in RIPA that are re-suspended in 9 M of urea. All proteins are normalized to tubulin in the soluble fraction and the relative level of each fraction at 0 h is set to 1. Mean ± SEM, n = 9 in (A-B), n = 3 in (D-F, J-L and N-P), n = 10~11 cells per time point from pooled results of 3 independent repeats in (H); statistic significance is determined by comparing to the level at 0 h in each test; *$p < 0.05$, ***$p < 0.001$; one-way ANOVA; ud, undetected. Scale bars: 5 μm.

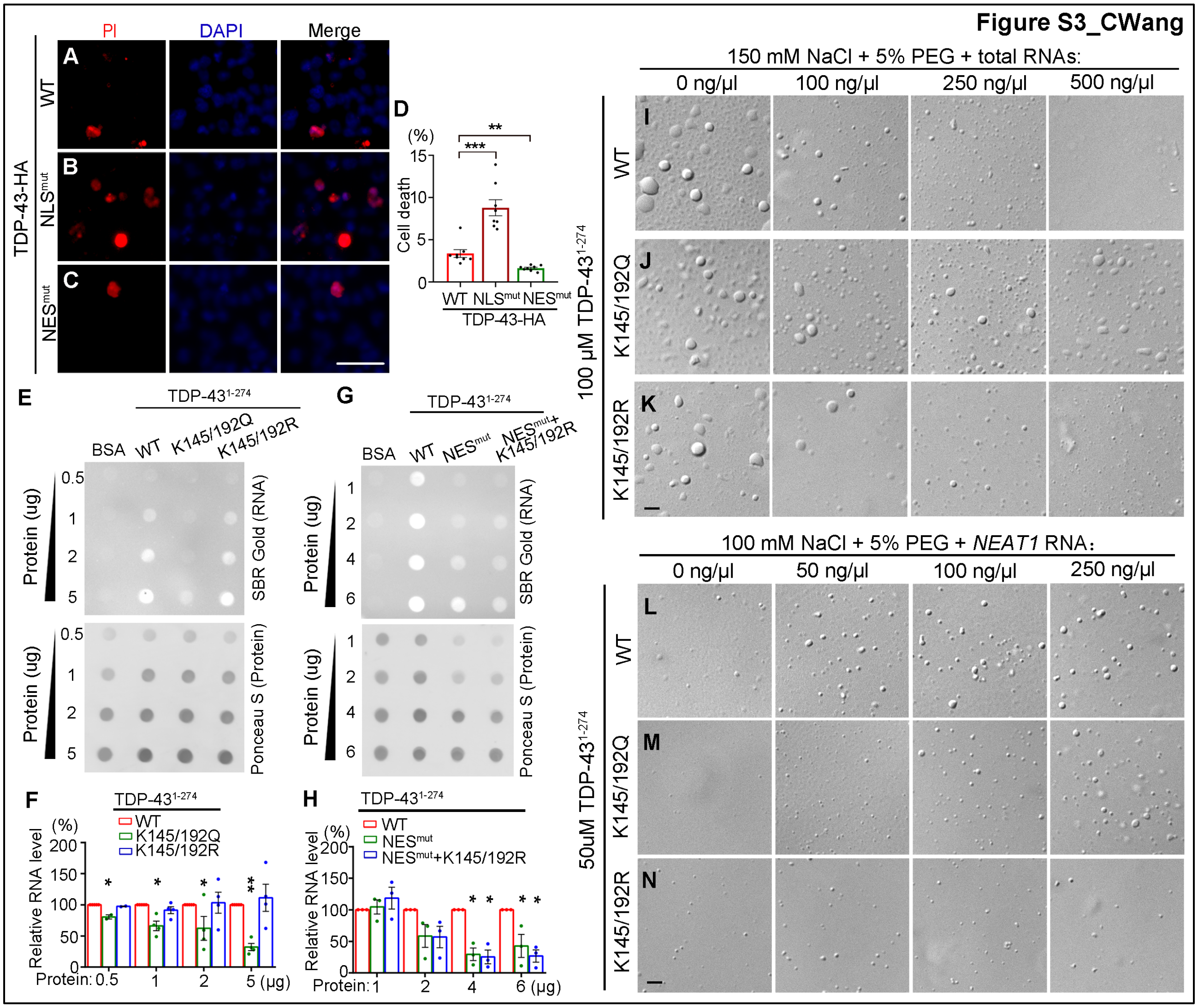


**Figure S3. Altered RNA binding in the NES$^{mut}$ and the K145/192 mutations may underlie the formation of TDP-43 droplets and the reduced cytotoxicity, related to Figure 3 and 5**

(**A-D**) Representative images (A-C) and quantification (D) of PI staining to evaluate the death of cells expressing WT or mutant TDP-43 as indicated. (**E-H**) Representative images (E, G) and quantifications (F, H) of the *in vitro* dot-blot assays evaluating the RNA binding affinity of WT or mutant TDP-43 as indicated. The bound RNA is visualized using the SYBR® Gold Nucleic Acid Gel Stain kit and the protein loaded is stained by Ponceau S. BSA is used as a negative binding control. (**I-K**) Suppression of TDP-43 LLPS by total RNA is impaired by the K145/192Q mutation but not enhanced by the K145/192R. (**L-N**) Promotion of TDP-43 LLPS by *NEAT1* RNA is drastically reduced by the K145/192R mutation. The concentrations of NaCl, TDP-43, PEG and RNA used in the *in vitro* assays are as indicated. Mean ± SEM; n = ~2000 cells each group from pooled results of 3 independent repeats in (D), n = 3~4 in (F, H); *$p$ < 0.05, **$p$ < 0.01, ***$p$ < 0.001; ns, not significant; Student $t$-test, by comparing to WT TDP-43 in each condition. Scale bars: 50 μm in (A-C), 2 μm in (I-N).

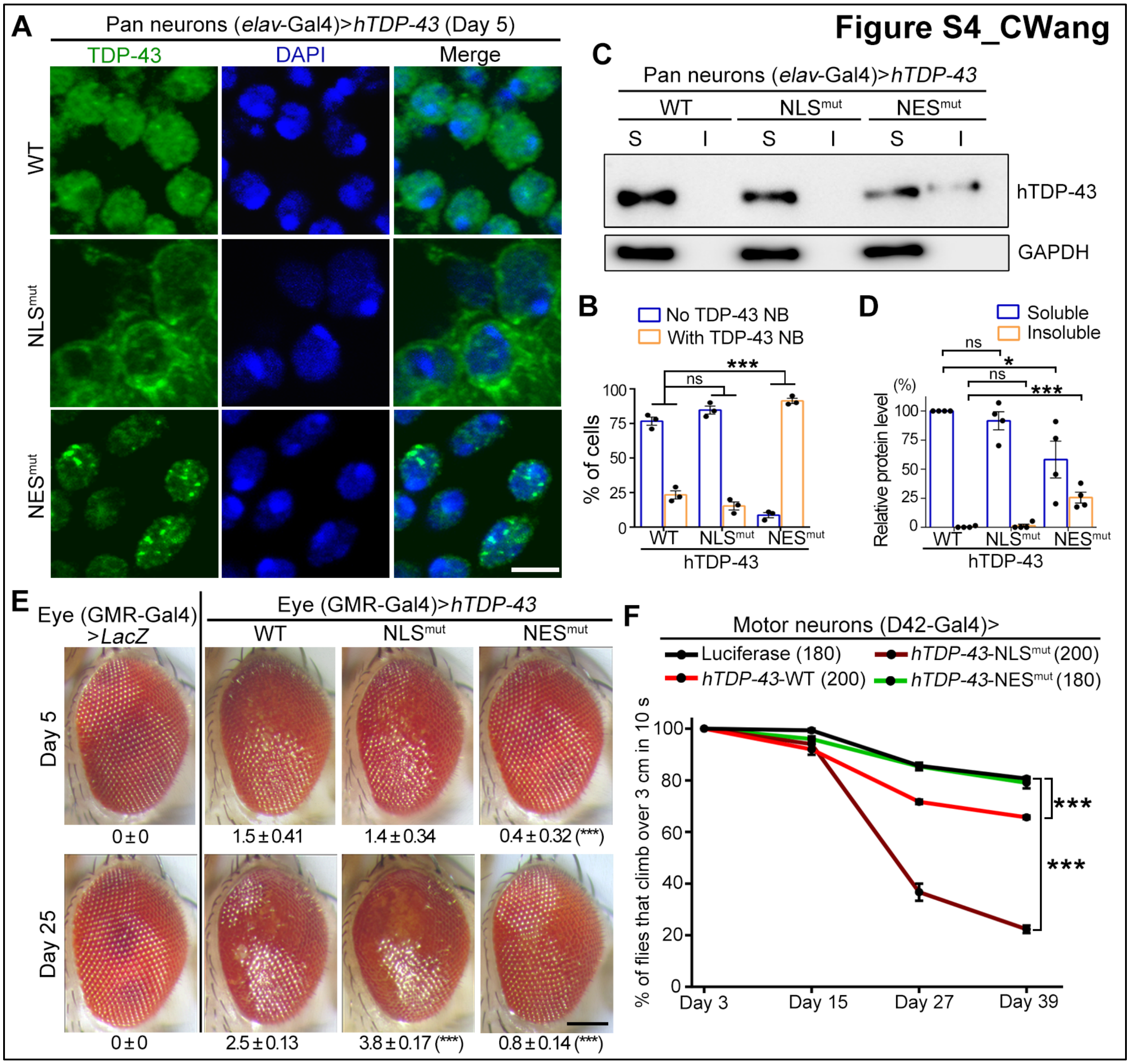


**Figure S4. TDP-43-NES$^{mut}$ forms NBs in fly neurons *in vivo* and that mitigates degenerative phenotypes in a *Drosophila* model of ALS, related to Figure 3**

(**A**) Representative confocal images of the fly central brain neurons (*elav*-Gal4) expressing WT or mutant *hTDP-43* as indicated. The whole-mount fly brains were dissected and immunostained for hTDP-43 (green) and DAPI (blue, to show the nucleus). (**B**) The average percentages of fly neurons with or without TDP-43 NBs are quantified. Mean ± SEM, n = ~100 neurons each group. (**C-D**) Representative image (C) and quantification (D) of Western blot analysis of WT or mutant hTDP-43 protein extracted from fly heads (Day 5) in RIPA-soluble (S) or RIPA-insoluble (I, resuspended in 9 M of urea) fractions. All protein levels are normalized to GAPDH. The relative level of soluble TDP-43-WT is set to 100%. Mean ± SEM, n = 4. (**E**) Representative z-stack images of the fly eye (driven by a GMR-Gal4 driver) expressing the WT or mutant hTDP-43 transgenes as indicated at Day 5 or Day 25. The degeneration severity is assessed by the rough surface of the compound eyes and loss of pigment cells. The average degeneration score (mean ± SEM) and the statistic significance (compared to hTDP-43-WT flies) of each group are indicated at the bottom. n = 8 eyes each group. (**F**) The climbing capability of the flies expressing the WT or mutant hTDP-43 in motor neurons (D42-Gal4) is evaluated as percentage of flies climbing over 3 cm in 10 seconds. The UAS-*luciferase* fly line is used as a control. Mean ± SEM, the number of flies tested for each genotype is shown. *$p < 0.05$, ***$p < 0.001$; ns, not significant; two-way ANOVA in (B, F), one-way ANOVA in (D-E). Scale bars: 2 μm in (A) and 100 μm in (E).

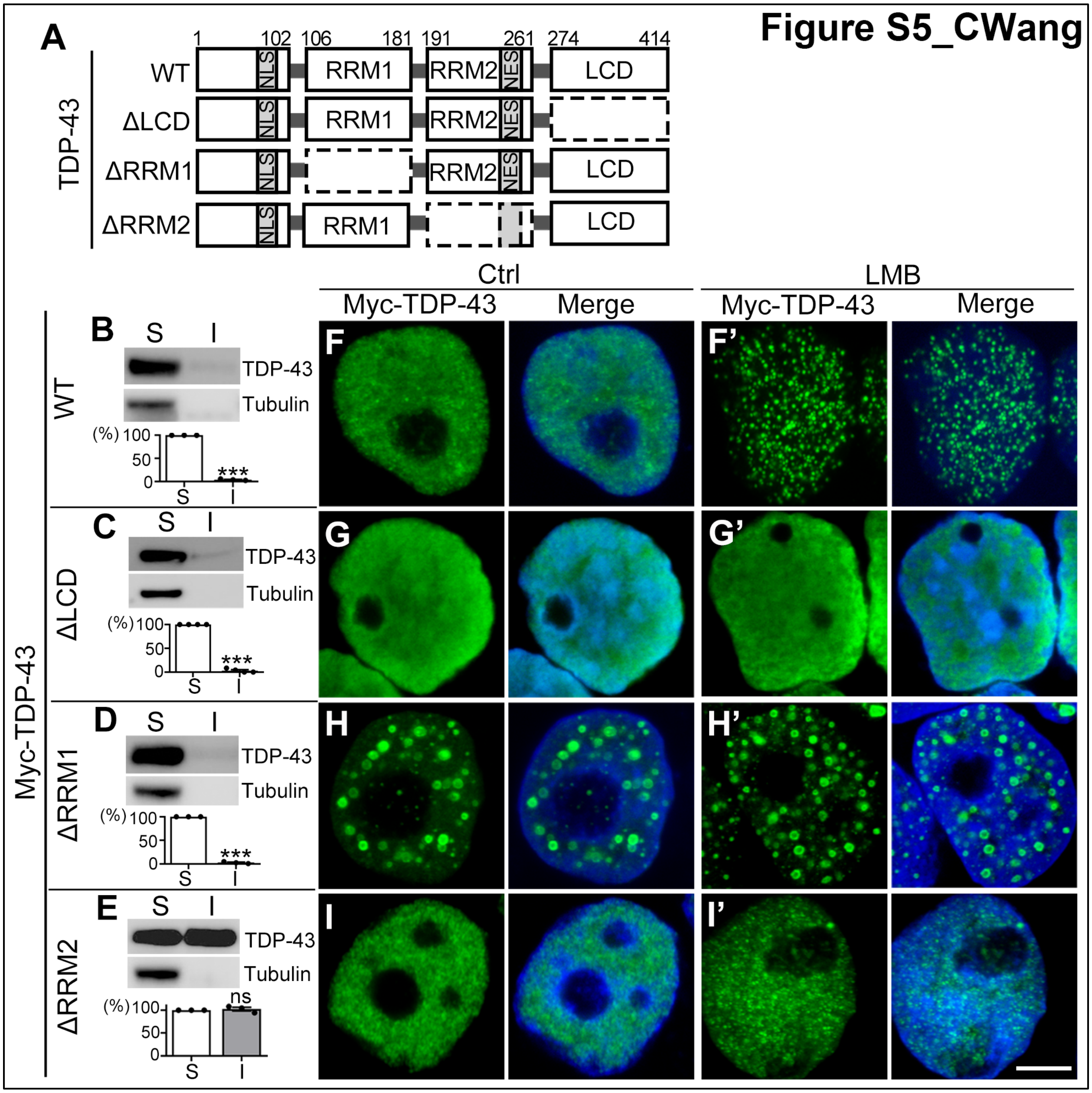


**Figure S5. The impact of the LCD and RRM domains on the assembly of TDP-43 NBs, related to Figure 4**

(**A**) A diagram showing the major functional domains and the truncated ΔLCD, ΔRRM1 and ΔRRM2 TDP-43. (**B-E**) Western blot analysis of TDP-43-WT (B), ΔLCD (C), ΔRRM1 (D) and ΔRRM2 (E) in the RIPA-soluble and RIPA-insoluble (resuspended in 9 M of urea) fractions. All TDP-43 proteins are normalized to tubulin in the soluble fraction and the relative level is quantified as percentage to the soluble fraction of each protein. Mean ± SEM, n = 3 independent repeats. (**F-I'**) Representative confocal images of TDP-43-WT, ΔLCD, ΔRRM1 and ΔRRM2 in 293T cells treated with the vehicle control ethanol (F-I) or LMB (F'-I'). Green, anti-Myc for Myc-TDP-43; blue, DAPI for DNA. Scale bar: 5 μm.

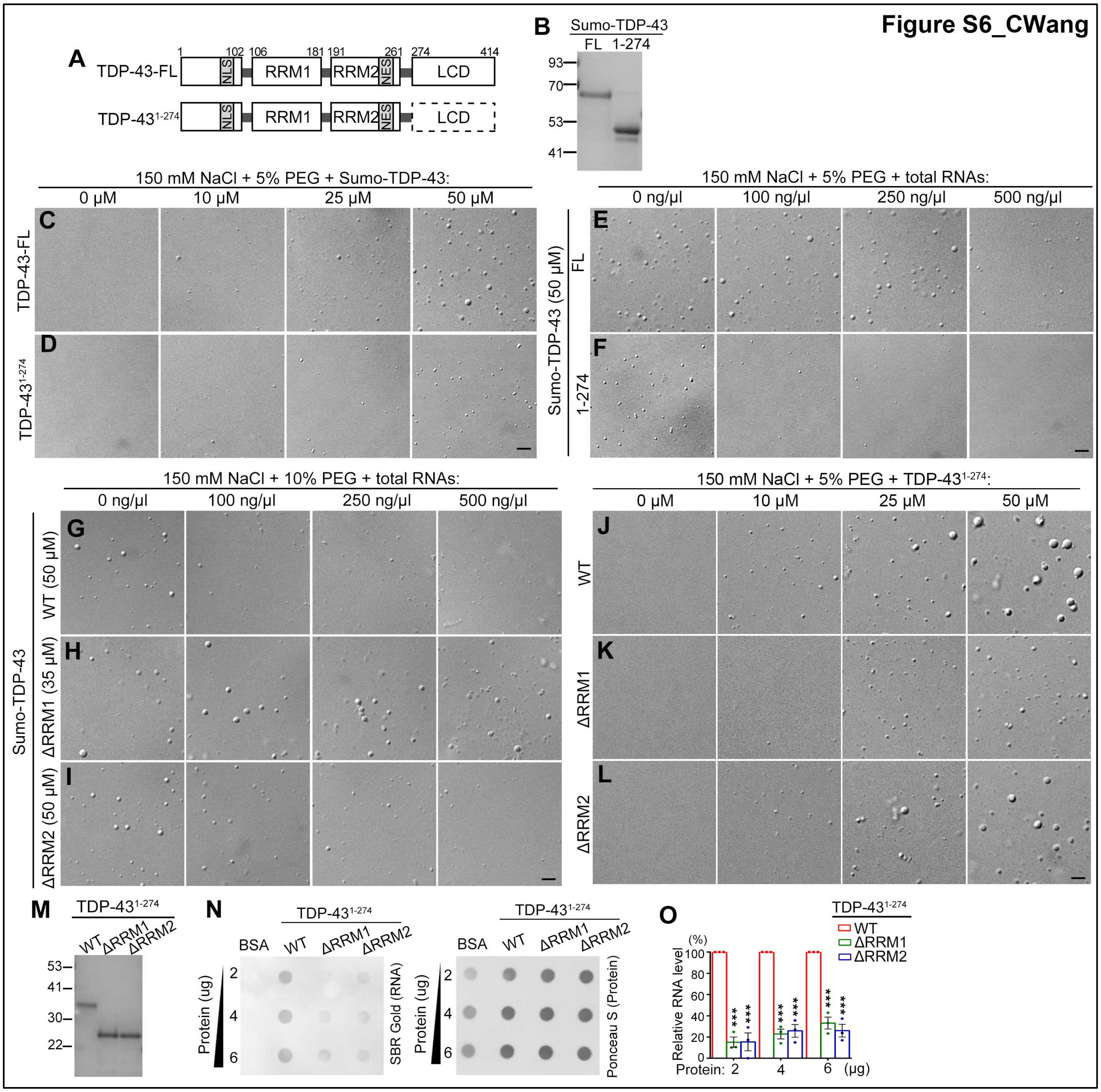


**Figure S6. Total RNAs suppress TDP-43 LLPS *in vitro*, related to Figure 5**

(**A**) A diagram showing the full-length (FL) and truncated TDP-43 protein without the LCD (TDP-43$^{1-274}$). (**B**) Coomassie blue staining confirming the purified FL and truncated sumo-TDP-43 proteins. (**C-D**) The *in vitro* LLPS of FL (C) and TDP-43$^{1-274}$ (D) is dose-dependent. (**E-F**) Suppression of the LLPS of the FL and the TDP-43$^{1-274}$ by total RNAs extracted from HeLa. (**G-I**) Suppression of the LLPS of the SUMO-tagged WT (G), ΔRRM1 (H) and ΔRRM2 (I) TDP-43 by total RNAs. Note that a different concentration in H is used for ΔRRM1 to reach a similar starting level of the size and number of TDP-43 droplets to WT and ΔRRM2 SUMO-TDP-43. (**J-L**) WT (J), ΔRRM1 (K) and ΔRRM2 (L) TDP-43$^{1-273}$ proteins forms LDs *in vitro* by LLPS in a dose-dependent manner. The concentrations of NaCl, the crowding agent PEG, TDP-43 protein and total RNAs used in the above assays are as indicated. (**M**) Coomassie blue staining confirming purified WT, ΔRRM1 and ΔRRM2 TDP-43$^{1-274}$ proteins used in the *in vitro* assays above and in Figure 5. (**N**) Representative images of the *in vitro* dot-blot assay confirming the reduced RNA binding affinity of the ΔRRM1 and ΔRRM2 TDP-43$^{1-273}$ compared to the WT. The bound RNA is visualized using the SYBR® Gold Nucleic Acid Gel Stain kit. Bovine serum albumin (BSA) is used as a negative binding control. (**O**) The RNA intensity is normalized to Ponceau S staining (protein loading control) and shown as the percentage to that of WT TDP-43$^{1-273}$ in (N). Mean ± SEM, n = 3; ***$p < 0.001$; one-way ANOVA. Scale bars: 2 μm.

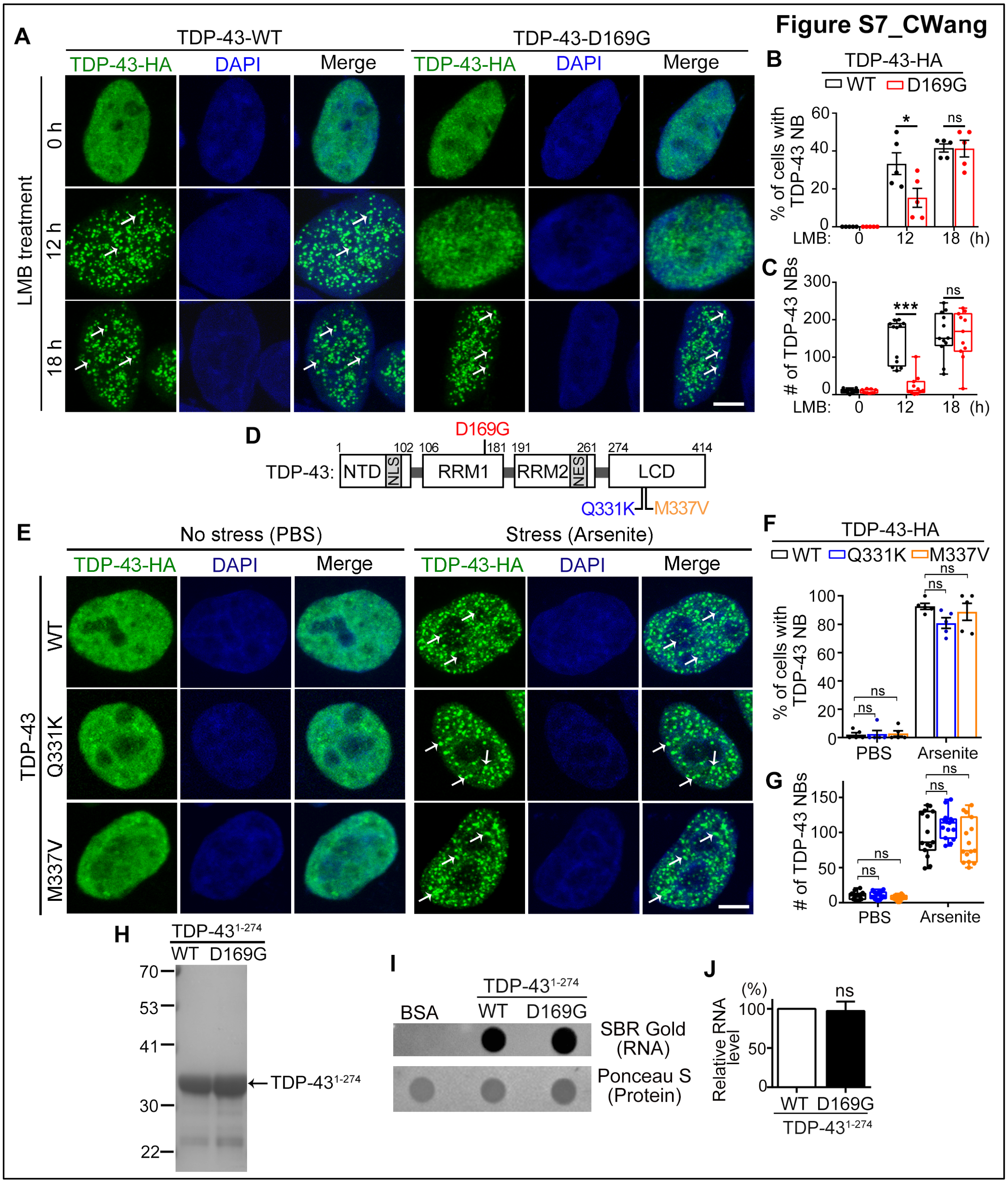


**Figure S7. A specific and unique defect in TDP-43 NB formation caused by the D169G mutation, related to Figure 6 and Figure 7**

(**A**) Representative confocal images of HeLa cells expressing with WT or D169G TDP-43 treated with LMB (25 nM) for indicated times. (**B-C**) Quantifications of the percentage of cells showing

TDP-43 NBs (B) and the average number of NBs per cell (C) at the end of the experiment in (A). LMB-induced NB formation was delayed in D169G TDP-43. (**D**) Diagrams showing WT TDP-43 and the mutations of D169G, Q331K and M337V in the LCD. (**E**) Representative confocal images of cells transfected with WT, Q331K or M337V TDP-43 in the absence (PBS) or presence of arsenite. Green, TDP-43-HA (anti-HA); Blue, DAPI staining for DNA (blue); arrows, TDP-43 NBs. (**F-G**) The percentage of cells with TDP-43 NBs (F) and the average count of TDP-43 NBs per cell (G) in (E) are quantified. (**H**) Coomassie blue staining confirming the purified WT and D169G TDP-43$^{1-274}$ proteins (arrow) used in the *in vitro* LLPS assays. (**I-J**) Representative images (I) and quantifications (J) of the *in vitro* dot-blot assay to evaluate the binding affinity of WT and D169G TDP-43$^{1-274}$ to total RNAs. Mean ± SEM, n = ~100 cells for each group in (B, F) and 11~15 cells per group in (C, G) of pooled results from 3 independent repeats, n = 3 in (J); *$p$ < 0.05, ***$p$ < 0.001; ns, not significant; Student's *t*-test in (B-C, J), one-way ANOVA in (F-G). Scale bars: 5 μm.

### Supplemental videos

**Video S1. A representative time-lapse video of *live* HeLa cells showing the formation of GFP-TDP-43 NBs upon the arsenite treatment**

**Video S2. A representative time-lapse video of GFP-TDP-43-GFP NBs in *live* HeLa cells before and after photobleaching**
